# Dissecting Succulence: Crassulacean Acid Metabolism and Hydraulic Capacitance are Independent Adaptations in *Clusia* Leaves

**DOI:** 10.1101/2022.03.30.486278

**Authors:** Alistair Leverett, Samantha Hartzell, Klaus Winter, Milton Garcia, Jorge Aranda, Aurelio Virgo, Abigail Smith, Paulina Focht, Adam Rasmussen-Arda, William G. T. Willats, Daniel Cowan-Turner, Anne M. Borland

## Abstract

- Succulence is found across the world as an adaptation to water-limited niches. The fleshy organs of succulent plants develop via enlarged photosynthetic chlorenchyma and/or achlorophyllous water storage hydrenchyma cells. The precise mechanism by which anatomical traits contribute to drought tolerance is unclear, as the effect of succulence is multifaceted. Large cells are believed to provide space for nocturnal storage of malic acid fixed by crassulacean acid metabolism (CAM), whilst also buffering water potentials by elevating hydraulic capacitance (C_FT_). Furthermore, the effect of CAM and elevated C_FT_ on growth and water conservation have not been compared, despite the assumption that these adaptations often occur together.
- We assessed the relationship between succulent anatomical adaptations, CAM and C_FT_, across the genus *Clusia*. In addition, we simulated the effects of CAM and C_FT_ on growth and water conservation during drought using the Photo3 model.
- Within *Clusia* leaves, CAM and C_FT_ are independent traits: CAM requires large palisade chlorenchyma cells, whereas hydrenchyma tissue governs interspecific differences in C_FT_. In addition, our model suggests that CAM supersedes C_FT_ as a means to maximise CO_2_ assimilation and minimise transpiration during drought.
- Our study challenges the assumption that CAM and C_FT_ are mutually dependent traits within succulent leaves.

## Introduction

The advent of global warming is expected to cause more frequent and severe drought events to ecosystems, worldwide (Choat *et al.*, 2018; Jiao *et al.*, 2021; Sheffield & Wood, 2008). To prepare for increased aridity, it is imperative that scientists develop a thorough understanding of the adaptations that plants currently employ to survive drought in nature (Borland *et al.*, 2015; Germon *et al.*, 2019; Trueba *et al.*, 2019; Yang *et al.*, 2020; Fradera-Soler *et al.*, 2021; Heyduk *et al.*, 2021). One adaptation that can be found across the world’s semi-arid ecosystems is succulence (Arakaki *et al.*, 2011; Jolly *et al.*, 2021; Merklinger *et al.*, 2021). Succulent plants can be recognised by their fleshy leaves and/or stems, which can store substantial volumes of water long after rainfall events have ended. Despite having convergently evolved across the globe, succulence is a complex syndrome that can develop in several different ways: water can be stored within large photosynthetic chlorenchyma cells, and/or in specialised achlorophyllous water-storage hydrenchyma cells (Fig. **1**) (Borland *et al.*, 2018; Heyduk, 2021; Males, 2017; Ogburn & Edwards, 2010). The consequence of succulence is multifaceted, often thought to be affecting both photosynthetic and hydraulic physiology. One key photosynthetic adaptation associated with succulence is crassulacean acid metabolism (CAM). In contrast to C_3_ photosynthesis, CAM plants assimilate the majority of their CO_2_ at night, temporarily fixing carbon into malic acid and storing it in the vacuole until the following day. By fixing CO_2_ at night, rather than during the day, CAM allows stomata to open when the atmosphere is cooler and more humid, which curtails transpirational water losses (Abraham *et al.*, 2020; Winter *et al.*, 2005). CAM requires large succulent chlorenchyma cells to provide sufficient space to store malic acid overnight (Nelson *et al.*, 2005; Barrera-Zambrano *et al.*, 2014; Males, 2018; Töpfer *et al.*, 2020). However, succulence also performs a further, hydraulic function, by elevating bulk hydraulic capacitance (C_FT_) (Borland *et al.*, 2018; Ogburn & Edwards, 2012). C_FT_ is defined as the ratio with which relative water content (RWC) and Ψ drop during dehydration, multiplied by the moles of stored water in a given leaf area (Equation 2). Organs with higher C_FT_ can lose more water without severe drops to Ψ, thereby avoiding hydraulic damage to vascular and mesophyll tissues (Brodribb *et al.*, 2016; Zhang *et al.*, 2016; Scoffoni *et al.*, 2017; John *et al.*, 2018). Therefore, succulence can help plants survive drought by aiding the CAM pathway which minimises water loss, and by elevating C_FT_, to buffer water potentials during dehydration.

**Fig. 1.**
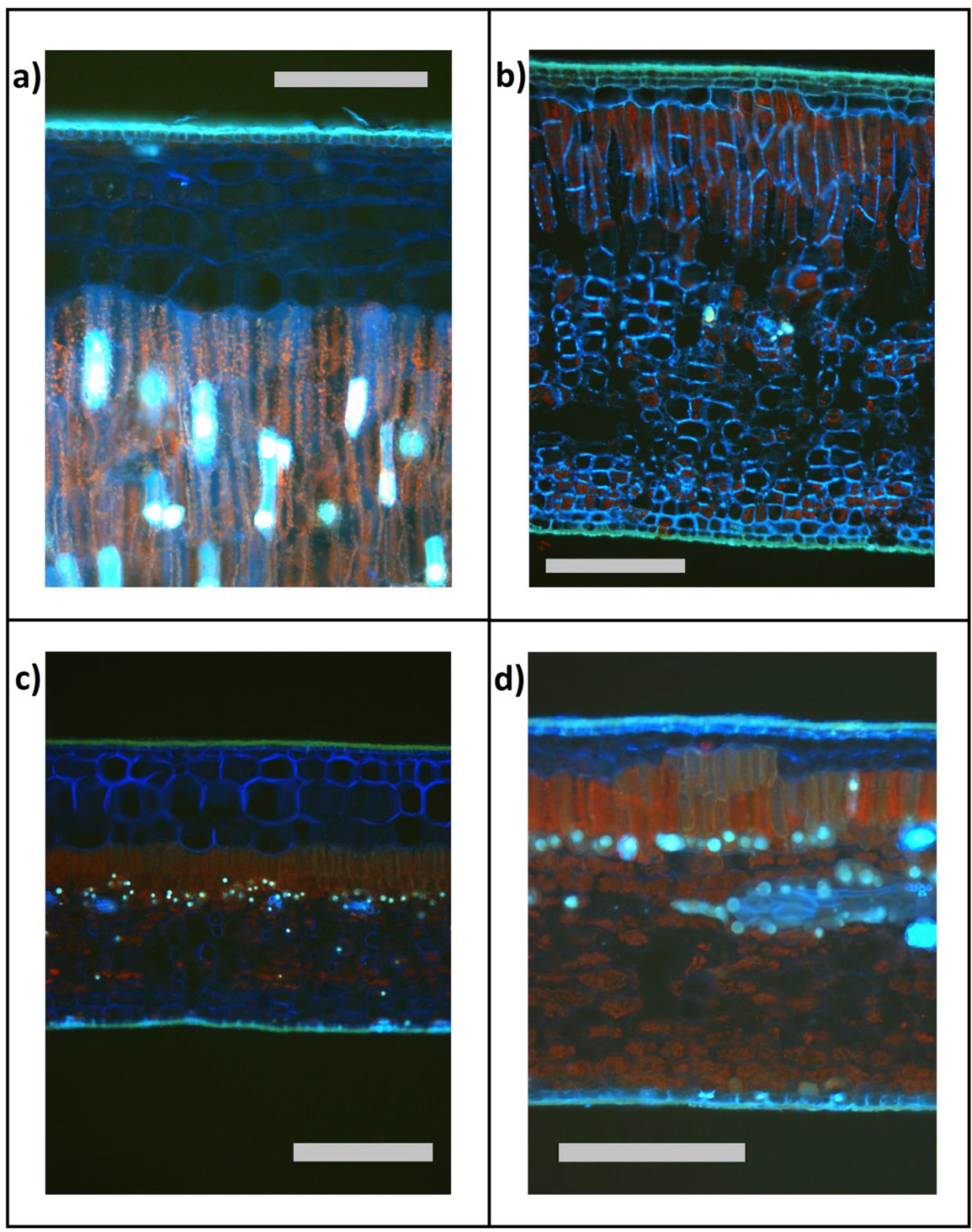
Anatomy of *Clusia* leaves, showing achlorophyllous hydrenchyma and photosynthetic chlorenchyma (comprised of palisade and spongy mesophyll layers). (a) Constitutive CAM species, *C. alata*, has deep hydrenchyma and deep chlorenchyma. (b) Constitutive CAM species, *C. rosea*, has deep chlorenchyma and shallow hydrenchyma. (c) Obligate C3 species, *C. multiflora*, has deep hydrenchyma and shallow chlorenchyma (d) Obligate C3 species, *C. grandiflora*, has shallow hydrenchyma and shallow chlorenchyma. For all images, scale bars represents 250 μm.

As CAM and a high C_FT_ are both believed to act as adaptations to drought, it is uncertain if one trait or both have driven the evolution of succulence. Strong CAM species often have large, tightly packed photosynthetic chlorenchyma, whereas most studies find little to no correlation between hydrenchyma depth and CAM expression (Barrera-Zambrano *et al.*, 2014; Males, 2018; Martin *et al.*, 2019). However, as CAM plants are typically found in water-limited niches, it is unclear if large chlorenchyma cells truly exist to provide vacuole space for malic acid, or if this anatomical adaptation is in fact a consequence of maximising C_FT_ to buffer plant water potentials during drought. A third possibility is that large chlorenchyma cells may have evolved to serve a dual function: providing space for malic acid storage and increasing C_FT_ (Edwards, 2019). Moreover, it is unclear if the presence of succulent hydrenchyma tissue can obviate the need for large chlorenchyma cells to provide C_FT_. If this were the case, it would indicate that large chlorenchyma cells are not playing a dual role, but are in fact a direct consequence of the evolution of strong CAM physiology. Building a robust physiological framework to describe the relationship between anatomy and C_FT_ is required to fully understand the role that succulence plays in facilitating CAM and address the potentially confounding effect that C_FT_ has on the relationship between large chlorenchyma cells and CAM. It is also unclear how CAM and C_FT_ interact to affect plant physiology during drought (Veste & Herppich, 2021). Both characteristics are thought to prevent plant Ψ from dropping, but it is not known if these adaptations function synergistically or if the presence of one obviates the need for the other. In short, whilst it is often assumed that CAM and elevated C_FT_ are found together, very little has been done to disentangle the effects that these adaptations have for plant physiology during drought. The use of modelling is ideal for this purpose, as it would allow CAM and C_FT_ to each be independently altered, whilst keeping other traits constant (Chomthong & Griffiths, 2020; Luo *et al.*, 2021).

To disentangle the roles of CAM and C_FT_ in succulent leaves, we analysed *Clusia*; a large phenotypically diverse genus of trees and hemiepiphytes from Central and South America and the Caribbean (Tinoco Ojanguren and Vázquez Yanes, 1983; Borland *et al.*, 1992; Holtum *et al.*, 2004; Leverett *et al.*, 2021; Luján *et al.*, 2021). Under well-watered conditions, *Clusia* leaves contain a diversity of photosynthetic phenotypes, ranging from obligate C_3_, to strong CAM (Barrera-Zambrano *et al.*, 2014). This range includes several species exhibiting C_3_-CAM intermediate phenotypes, where flux through the CAM cycle represents only a small to moderate fraction of total carbon assimilation (Winter, 2019; Tay, *et al.*, 2021). In addition, many *Clusia* species exhibit facultative CAM phenotypes, meaning they can increase metabolic flux through the CAM cycle in response to drought (Leverett *et al.*, 2021; Winter & Holtum, 2014). Furthermore, the leaves of *Clusia* exhibit a high degree of anatomical diversity (Luján *et al.*, 2021). Across a well characterised glasshouse collection of *Clusia*, CAM was found to correlate with enlarged palisade chlorenchyma cells (Barrera-Zambrano *et al.*, 2014). Moreover, across this *Clusia* collection there is substantial interspecific variation in adaxial hydrenchyma cell size and tissue depth, which is independent of CAM (Barrera-Zambrano *et al.*, 2014; Borland *et al.*, 2018). High photosynthetic and anatomical variation within closely related species makes it possible to conduct comparative analyses (Scoffoni *et al.*, 2016), which we used to dissect the roles of hydrenchyma from those of large palisade chlorenchyma cells. We interrogated the same *Clusia* collection as Barrera-Zambrano *et al.* (2014), to assess whether large palisade chlorenchyma cells correlate with C_FT_, as they do with CAM. Furthermore, we investigated saturated water content (SWC), cell wall characteristics and osmotic properties of leaves to understand the physiological traits that lead to interspecific differences in C_FT_. Finally, we parameterised the recently developed Photo3 model (Hartzell *et al.*, 2018; 2021) with *Clusia* data, to establish the roles that CAM and C_FT_ each play during drought. Together, these analyses deconstruct the assumption that CAM and C_FT_ are mutually dependent traits within succulent leaves.

## Materials and Methods

### Plant Growth Conditions

A previously characterised glasshouse collection of 11 *Clusia* species was used for comparative analyses. Plants aged 3-6 years (approx. 60-100 cm tall) were grown in 3:1, (v/v) compost-sand mixture (John Innes No. 2, Sinclair Horticulture Ltd, Lincoln, UK), in 10 L pots. Plants aged 3-6 years (approx. 60-100 cm tall) were grown in 3:1, (v/v) compost-sand mixture (John Innes No. 2, Sinclair Horticulture Ltd, Lincoln, UK), in 10 L pots. Plants were lit with sunlight plus photosynthetic LED lights (Attis 5 LED plant growth light, PhytoLux) to ensure that a 12 h photoperiod was maintained throughout the year. Plants were watered every two days and glasshouse temperatures were maintained at 25 and 23 °C during the day and night, respectively.

*Clusia alata* (constitutive CAM) and *C. tocuchensis* (obligate C_3_) plants were used for leaf dissections (described below). These plants were propagated from cuttings, as described in Barrera-Zambrano *et al.*, (2014). The 3–4-year-old plants were then transferred to a plant growth chamber (SANYO Fitotron) and allowed to acclimatise for 3 weeks (12-hour light period, day/night temp 25/19 °C). Plants received approx. 500 μM m^-2^ s^-1^ PFD at leaf height and were watered every two days.

### Gas Exchange

Published gas exchange data was used as a quantitative estimate of investment in CAM (Barrera-Zambrano *et al.*, 2014; Leverett *et al.*, 2021). CAM was estimated by measuring the percentage of diel CO_2_ assimilation occurring at night under well-watered conditions (CAM_ww_) and after 9 days of drought (CAM_d_).

Stomatal conductance to H_2_O, for *C. alata* and *C. tocuchensis*, was recorded using a LI- 6400XT infrared gas analyser (LiCOR, Lincoln, NE, USA). Leaves were sealed into a leaf chamber fluorometer cuvette, set to track external light conditions with humidity maintained around 60 % and data was logged every 15 minutes. Three biological replicates were made for each species, and one representative graph is included to show stomatal conductance for each species.

### Anatomical Measurements

Estimates of palisade, spongy mesophyll and hydrenchyma cell sizes were taken from Barrera-Zambrano *et al.*, (2014) and Borland *et al.*, (2018). Estimates of tissue depth were generated by hand sectioning leaf lamina, midway along the proximal-distal axis. Sectioned material was imaged using a Leitz Diaplan and a GXCAM HiChrome-S camera (GT Vision Ltd), or a with a Leica DM6B microscope and a Leica DFC9000 sCMOS camera (Leica microsystems).

### Pressure-Volume Curves

Pressure-volume curve data for 11 *Clusia* species were taken from Leverett *et al.*, (2021). These data were used to plot leaf water potential (Ψ_L_) or turgor pressure (P) against relative water content (RWC), where

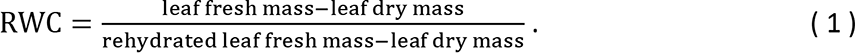

Bulk hydraulic capacitance (CFT) was calculated as

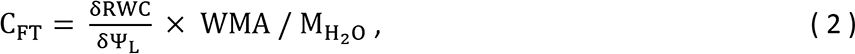

where WMA is leaf water mass per leaf area and MH2O is the molar mass of water (Beadle *et al.*,

1985; Blackman & Brodribb, 2011; Males & Griffiths, 2018). The bulk modulus of cell wall elasticity (ε) was calculated as

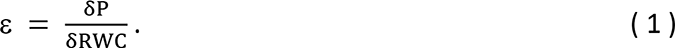

For pressure-volume curves, n = 6. Species mean C_FT_ was compared to published cell size data (Barrera-Zambrano *et al.*, 2014; Borland *et al.*, 2018). In addition, we sought to develop a larger data set, with which to conduct more robust statistical analyses. To this end, we built a linear mixed effect model to estimate C_FT_ based on SWC (fixed effect) and species (random effect). Using this model, C_FT_ could be estimated, alongside tissue depth for each leaf. Modelled C_FT_ correlated tightly with measurements from pressure-volume curves (Fig. **S1**: slope = 1.07, R^2^ = 0.95, p < 0.0001). These leaf-level data were used to construct linear mixed effect models to assess the relationships between tissue depth and C_FT_ (n = 12).

### Dissection and measurement of relative water content (RWC)

Hydrenchyma and chlorenchyma tissues were dissected in *C. alata* and *C. tocuchensis* since these species contain deep hydrenchyma, making them amenable to dissection (Fig. **S2**). Hydrenchyma tissue was removed from chlorenchyma, using the blunt edge of a razor. Dissected material was inspected, using a Leica DM6B microscope (Fig. **S3**) to ensure that clean separation was possible. Each dissection took < 1 s.

Relative water content (RWC) was measured every 4 hours for 24 h by quickly weighing dissected material which was then placed in deionised water to rehydrate for 8 h. Chlorenchyma tissue often sank, indicating that internal air space was filling with water and altering the validity of these data (Arndt *et al.*, 2015). Consequently, only the RWC of the hydrenchyma could be analysed and was calculated according to equation 1.

### Structural Carbohydrate Profiling

Comprehensive Microarray Polymer Profiling (CoMPP) was performed according the methodology of Moller *et al.*, (2007), with slight adjustments. Tissue was ground with pestle and mortar in liquid nitrogen, then mixed with 70% ethanol (1 mL). Samples were centrifuged (12,000 g, 10 minutes) and the supernatant discarded. This was repeated with methanol:chloroform (1:1 ratio, 1 mL), then acetone and pellets were left to dry overnight to form an alcohol insoluble residue (AIR).

CDTA and NaOH were used to sequentially extract polysaccharides from AIR samples. A non-contact microarray printer (Arrayjet, Roslin, UK) was used to print samples onto a nitrocellulose membrane, and these arrays were probed (each array probed separately) with a panel of antibodies with specificities for different polysaccharide epitopes. Relative spot signal intensities arising from antibody binding were determined using microarray analysis software (ArrayPro- Analyser, Media Cybernetics, Rockville, USA) before creating a heatmap based on mean signal intensities.

### Titratable Acidity and Osmometry

Leaves were harvested 15 minutes before dawn and before dusk for *C. alata* and *C. tocuchensis*. A 2-3 cm^2^ rectangle of leaf lamina was cut, midway along the proximal-distal axis of the leaf. The hydrenchyma and chlorenchyma layers were dissected, and these tissue layers were immediately frozen in liquid nitrogen. Frozen samples were crushed using a tissue lyser (Quiagen), incubated in 80 % methanol (Fisher) for 45 minutes and centrifuged at 14000g for 10 minutes. The supernatant was titrated against 0.5 Mol NaOH (Sigma), using phenolphthalein (Sigma) as an indicator, to determine H^+^ concentration.

Osmolality was measured for the same dawn-dusk samples using an Osmomat 030 cryoscopic osmometer (Gonotec). Lysed leaf tissue was vortexed in 200 μL deionised water, and 20 μL of this extract was placed into the osmometer to determine osmolality (Osmol kg^-1^). The osmotic potential was calculated according to the van’t Hoff equation.

### Statistics

Statistics were performed with R, v.3.6.0. Linear mixed effect models were built with the package ‘nlme’.

Pressure-volume curve parameters that are absolute (i.e. units are expressed on a per leaf area basis) were compared with absolute anatomical estimates, such as tissue depth or cell size. Pressure-volume curve parameters that are relative were compared with relative anatomical estimates, i.e. percentage of leaf depth comprised of hydrenchyma.

### Modelling of CAM and Hydraulic Capacitance

The previously developed Photo3 model, which uses recent advances in CAM modelling (Bartlett *et al.*, 2014; Hartzell *et al.*, 2015) to represent both C_3_ and CAM photosynthesis (Hartzell *et al.*, 2018) was parameterised with published photosynthetic traits from *Clusia*. In addition, the model was parameterised with the maximum and minimum C_FT_ values recorded in this study (*C. alata* and *C. grandiflora*, respectively), plus additional data collected for this purpose (Supplementary File 1). The model was run using climactic data from Gamboa, Panama to simulate both static soil moisture drought and dry-down conditions (Supplementary File 1). Climatic data from Gamboa was used because many species of *Clusia* grow in this location.

## Results

### Crassulacean Acid Metabolism (CAM) and Bulk Hydraulic Capacitance (CFT) are Independent in Clusia

The leaves of *Clusia* are comprised of photosynthetic palisade and spongy mesophyll layers (collectively the chlorenchyma) and achlorophyllous hydrenchyma (Fig. **1**). Hydrenchyma cell size/tissue depth varied independently of palisade and spongy mesophyll cell size/tissue depth (Fig. **S4**), hence the relative contribution of each tissue layer to CAM and C_FT_ could be assessed. Quantitative estimates of CAM expression used in this study were taken from Barrera Zambrano *et al.*, (2014) and Leverett *et al.*, (2021), who calculated the percentage of diel net CO_2_ assimilation occurring at night, in well-watered conditions and following 9 days of drought (CAM_ww_ and CAM_d_, respectively). Leaf water mass per area (WMA) correlated with both CAM_ww_ and CAM_d_, across 11 *Clusia* species (**Fig. 2a,b**). However, despite having more water, the leaves of CAM species did not exhibit elevated C_FT_: neither CAM_ww_ nor CAM_d_ correlated with C_FT_ (**Fig. 2c,d**). To understand why CAM and C_FT_ were independent, we tested the relationship between anatomical traits and C_FT_. A linear relationship was found between hydrenchyma cell size and C_FT_, but no such trend was observed for the cell sizes of other tissue layers (**Fig. 3a-c**). A similar conclusion was also drawn when tissue depth was analysed, in the place of cell size (**Fig. 3d-f**). A linear mixed effect model found that hydrenchyma depth significantly correlated with C_FT_, whereas the depth of palisade or spongy mesophyll did not. When this model was refined to remove spongy mesophyll, the conclusion did not change: the hydrenchyma depth significantly correlated with C_FT_ but palisade depth did not. Taken together, these data show that hydrenchyma, and not the palisade or spongy mesophyll, contributes to intraspecific variation in C_FT_, in *Clusia* leaves.

**Fig. 2.**
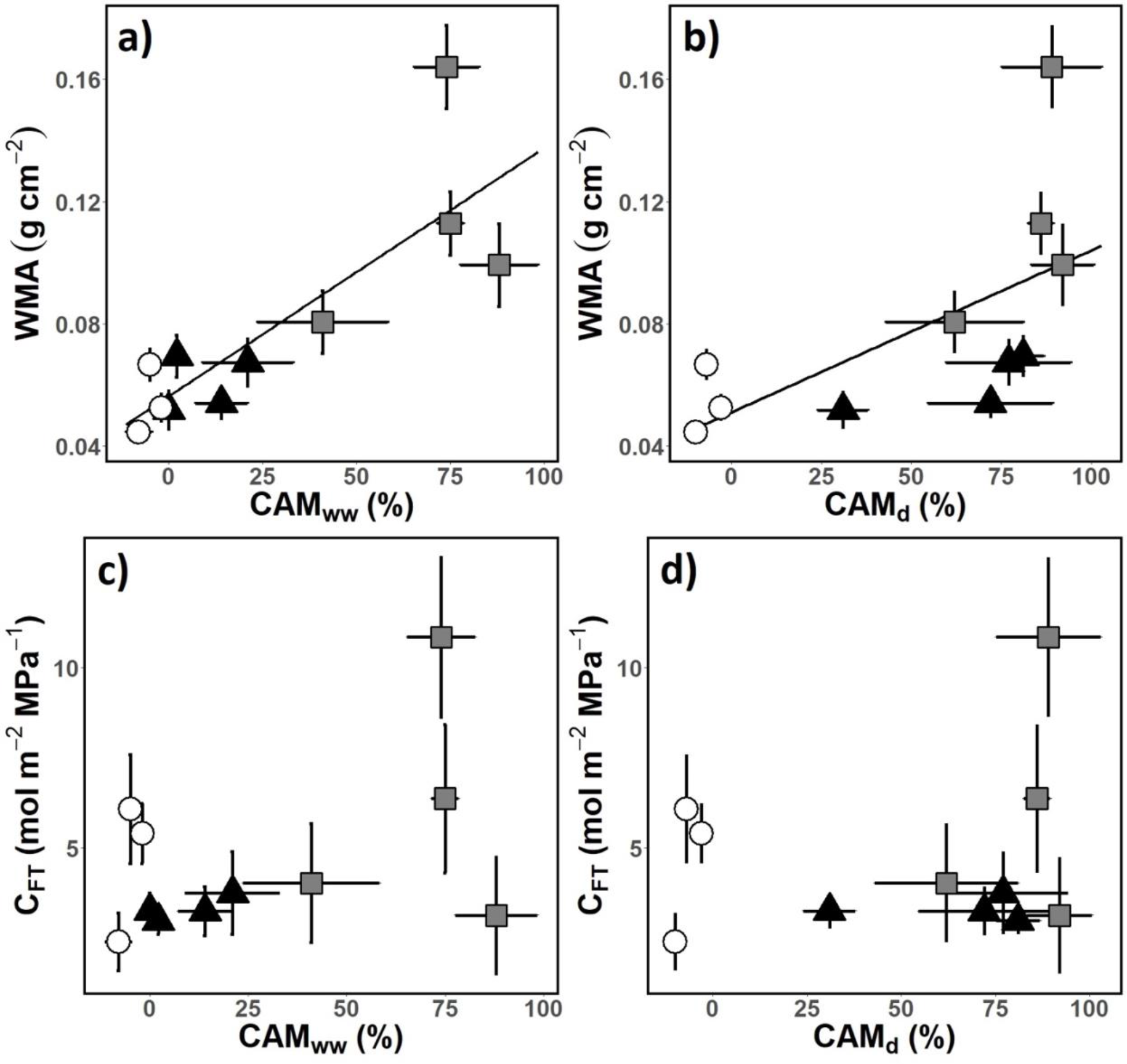
Species with Crassulacean acid metabolism have more water in leaves, but do not have elevated bulk hydraulic capacitance (C_FT_), in *Clusia*. Quantitative estimates of CAM expression were calculated as the percentage of diel CO_2_ assimilation occurring at night, in well-watered conditions and following 9 days of drought (CAM_ww_ and CAM_d_, respectively). (a) Across 11 species of *Clusia*, CAM_ww_ correlated with water mass per area (WMA) (linear regression: R_2_ = 0.69; p = 0.001) (b) CAM_d_ also correlated with WMA (linear regression: R^2^ = 0.38; p = 0.04). (c) CAM_ww_ did not correlate with C_FT_ (linear regression: p = 0.18). (d) CAM_d_ did not correlate with C_FT_ (linear regression: p = 0.63). White circles = obligate C_3_ species, black triangles = C_3_-CAM intermediate species and grey squares = constitutive CAM species. For each species, n = 6, error bars represent ± 1 standard deviation.

**Fig. 3.**
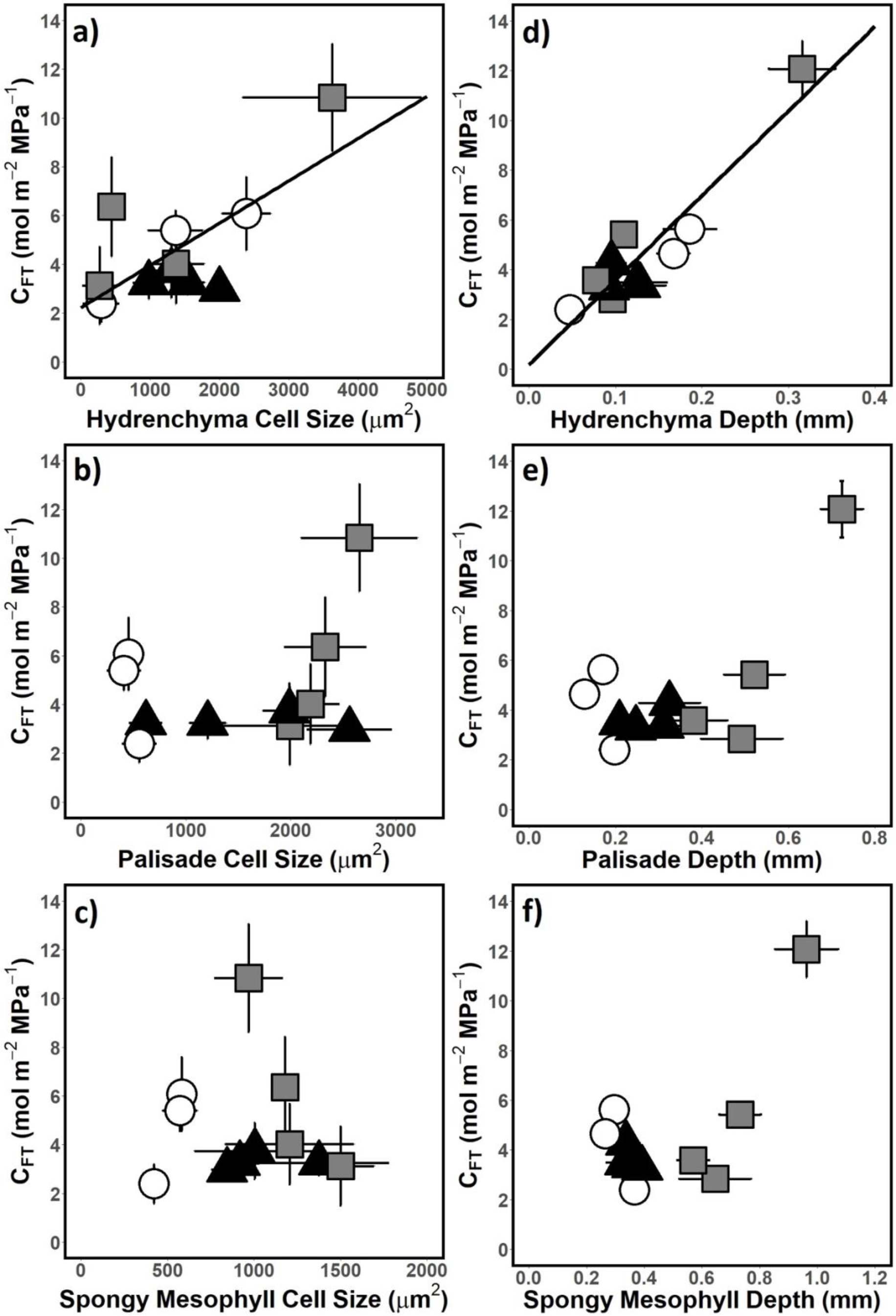
Hydrenchyma, and not the palisade or spongy mesophyll is responsible for interspecific variation in bulk hydraulic capacitance (C_FT_). (a) Hydrenchyma cell size positively correlated with C_FT_ (linear regression: R^2^ = 0.51; p = 0.01). Neither (b) palisade, nor (c) and spongy mesophyll cell size correlated with C_FT_ (linear regression: p = 0.37 and p = 0.85, respectively). (d-f) A linear mixed effect model was built to describe the relationship between tissue depths and C_FT_. This model found (d) a significant linear relationship between hydrenchyma depth and C_FT_ (p = 0.002) but no such relationship between (e) palisade depth and C_FT_ (p = 0.33) or (f) spongy mesophyll depth and C_FT_ (0.93). When this linear mixed effect model was refined by removing spongy mesophyll depth as a fixed effect, a likelihood ratio test found no significant difference (likelihood ratio = 3.27, p = 0.66) and the overall conclusion was unchanged: hydrenchyma depth significantly predicted C_FT_ (0.008) whereas palisade depth did not (0.2). When this second linear mixed effect model was refined by removing palisade depth as a fixed effect, a likelihood ratio test found no significant difference (likelihood ratio = 3.00, p = 0.56). For cell size data (a-c), linear regressions were built using species averages. For tissue depth data, linear mixed effect models were built using leaf-level data (n = 12). White circles = obligate C_3_ species, black triangles = C_3_-CAM intermediate species and grey squares = constitutive CAM species. Error bars represent ± 1 standard deviation.

We explored the relationship between CAM and saturated water content (SWC), as the latter has been shown to be predictive of capacitance in succulent taxa (Ogburn and Edwards, 2012). SWC did not correlate with either CAM_ww_ or CAM_d_ (**Fig. 4a,b**). Furthermore, SWC did not correlate with leaf thickness or WMA, when data from all 11 *Clusia* species were considered together (**Fig. 4c,d**). However, there was a clear linear relationship between WMA and SWC when only comparing species with C_3_ or C_3_-CAM photosynthetic physiologies (dashed lines, **Fig. 4c,d**). Likewise, a linear relationship existed between the WMA and SWC values for the strong CAM species (solid line, **Fig. 4c,d**). The lack of a significant relationship between SWC and WMA in the total data set was the result of the strong CAM species having higher WMA without a corresponding higher SWC (i.e. the gap between dashed and solid lines, **Fig. 4c,d**). This occurred because strong CAM species had higher leaf dry mass per area (LMA) (**Fig. 4e**). Taken together, these data show that higher WMA causes leaves to have higher SWC, unless this high WMA is associated with the presence of a strong CAM phenotype. Put differently, leaves of strong CAM species have more water, but do not exhibit high SWC; the physiological characteristic typically associated with high capacitance (Ogburn and Edwards, 2012).

**Fig. 4.**
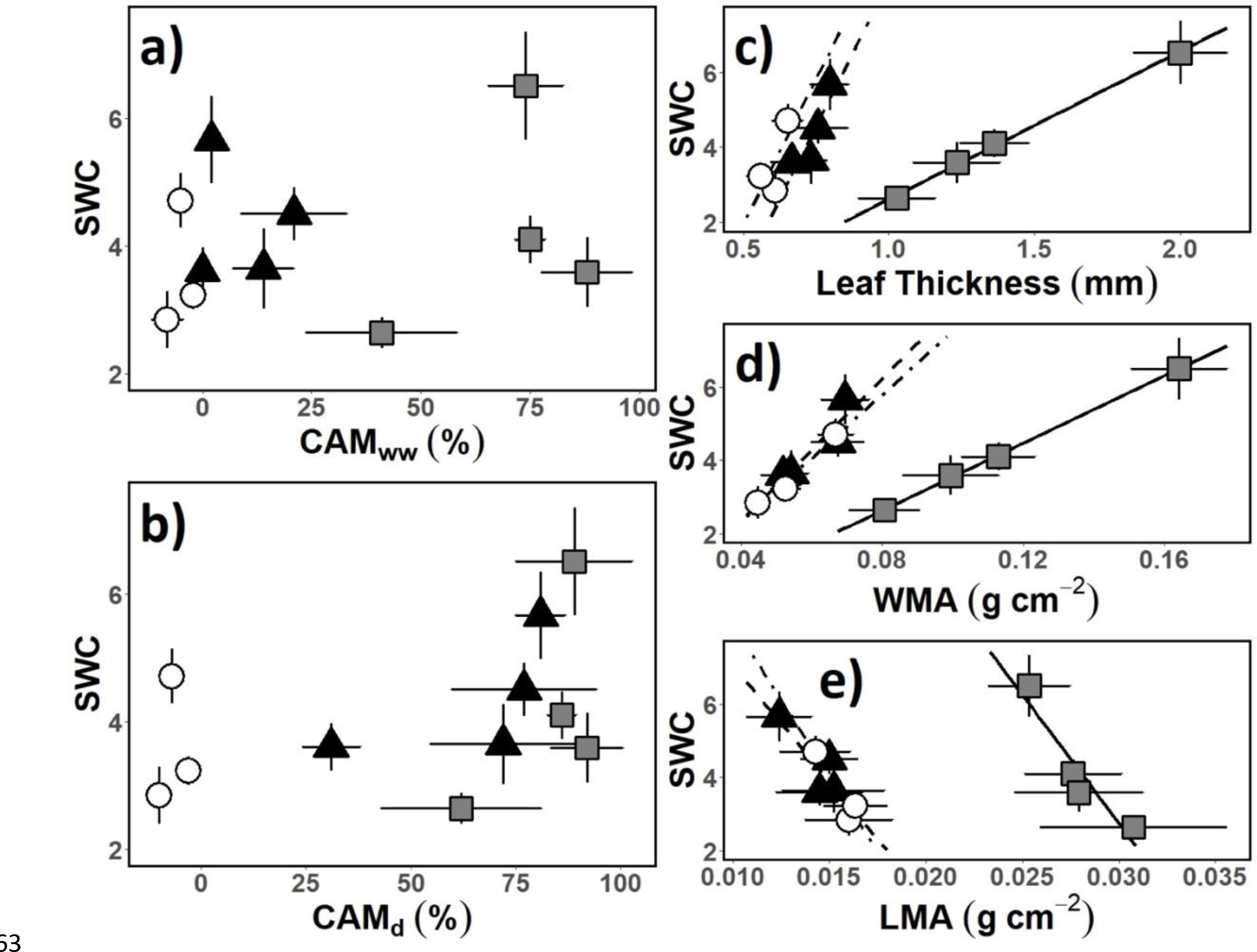
Crassulacean acid metabolism (CAM) and saturated water content (SWC) are independent in *Clusia* leaves. Quantitative estimates of CAM expression were calculated as the percentage of diel CO_2_ assimilation occurring at night, in well-watered conditions and following 9 days of drought (CAM_ww_ and CAM_d_, respectively). (a) Across 11 species of *Clusia* CAM_ww_ did not correlate with SWC (linear regression: p = 0.54). (b) CAM_d_ did not correlate with SWC (linear regression: p = 0.21). (c) Across 11 species of *Clusia* leaf thickness did not correlate with SWC. However, a clear linear relationship was seen when considering species with one photosynthetic physiotype: when the obligate C3 species (dot-dashed line), the C3-CAM species (dashed line) or the constitutive CAM species (solid line) were considered alone, there was a linear relationship between leaf thickness and SWC. Likewise, across 11 species, water mass per area (WMA) did not correlate with SWC. However, a clear linear relationship existed between WMA and SWC when considering only species with the same photosynthetic physiotype. (e) Leaf dry mass per area (LMA) did not correlate with SWC across 11 species of *Clusia*. However, a linear relationship existed between LMA and SWC when considering only species with the same photosynthetic physiotype. For all graphs, n = 12, white circles = obligate C3 species, black triangles = C3-CAM intermediate species and grey squares = constitutive CAM species. Error bars represent ± 1 standard deviation.

### Elastic Cell Walls Confer CFT in the Hydrenchyma

We sought to identify the properties of the hydrenchyma that confer increased C_FT_. One trait thought to contribute to C_FT_ is elastic cell walls, as they allow cells to readily deform and mobilise water stores. We estimated the bulk modulus of cell wall elasticity (ε: a larger value represents cell walls that are highly rigid). Across 11 species of *Clusia* C_FT_ negatively correlated with ε, which in turn negatively correlated with the percentage of leaf depth made up of hydrenchyma (**Fig. 5a,b**). Taken together, these data suggest that hydrenchyma cell walls are highly elastic and that this elasticity contributes to elevating C_FT_.

**Fig. 5.**
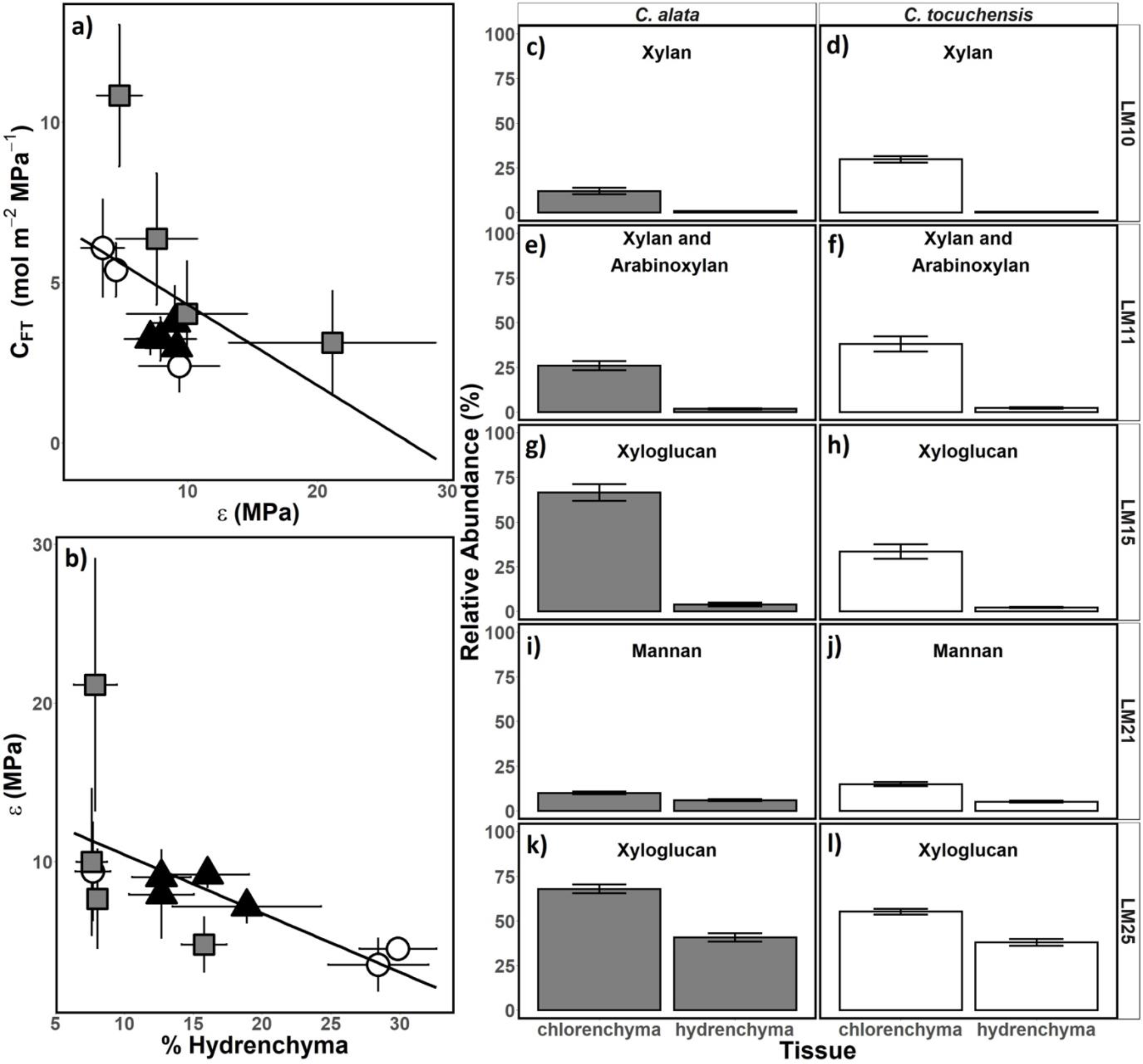
Elastic cell walls in the hydrenchyma confer elevated bulk hydraulic capacitance (C_FT_), in *Clusia*. (a) Across 11 species of *Clusia*, the bulk modulus of cell wall elasticity (ε) negatively correlated with C_FT_ (linear regression: R^2^ = 0.31; p < 0.001). (b) Across 11 species of *Clusia*, ε negatively correlated with the percentage of leaf depth comprised of hydrenchyma (% hydrenchyma) (linear regression: R^2^ = 0.39; p = 0.03). (c-g) Hemicellulose contents were lower in the hydrenchyma in the constitutive CAM species, *C. alata* (two tailed t-test: p < 0.001 for all graphs). (h-l) Hemicellulose contents were lower in the hydrenchyma in the obligate C_3_ species, *C. tocuchensis* (two tailed t-test: p < 0.001 for all graphs). Conclusions were unchanged when a Bonferroni-Hochberg adjustment was applied to p-values to account for multiple comparisons.

To examine how the composition of hydrenchyma cell walls impacts ε, the structural carbohydrate composition of cell walls was analysed. Hydrenchyma was dissected from chlorenchyma in both *C. tocuchensis* (obligate C_3_) and *C. alata* (constitutive CAM), and cell wall composition assessed using antibodies raised against 10 common polysaccharide and glycoprotein motifs. In both species, hemicelluloses (xylan, mannan and xyloglucan epitopes) were considerably less abundant in the hydrenchyma, compared with the chlorenchyma (Fig.5c- l). Together, these data suggest that low ε in the hydrenchyma is likely the result of this tissue having reduced structural reinforcement from hemicelluloses. In addition, in *C. alata* pectins were more abundant in the hydrenchyma than the chlorenchyma, although this pattern was not observed in *C. tocuchensis* (Fig. **S5**).

### High Osmotic Potentials Confer CFT in the Hydrenchyma

Whilst high ε values allow hydrenchyma cells to deform to mobilise water, movement of water into the chlorenchyma can only occur if an osmotic gradient exists between the tissues. We measured the osmotic potential (π) of dissected hydrenchyma and chlorenchyma tissues in *C. alata* and *C. tocuchensis*. Values of π were higher (less negative) in the hydrenchyma than the chlorenchyma for both species (**Fig. 6**). In the constitutive CAM species, *C. alata*, there was a significant change in chlorenchyma π between dawn and dusk, due to fluctuations in acid content in this tissue (**Fig. 6a**). However, π remained higher in the hydrenchyma than the chlorenchyma at both dawn and dusk (**Fig. 6b**). In the obligate C_3_ species, *C. tocuchensis*, chlorenchyma π did not differ with time. As with *C. alata*, π was higher in the hydrenchyma than the chlorenchyma at both dawn and dusk (**Fig. 6**). Together, these data show that an osmotic gradient exists between the hydrenchyma and the chlorenchyma. In addition, these data show that fluctuations in acid contents from CAM do not eliminate this osmotic gradient.

**Fig. 6.**
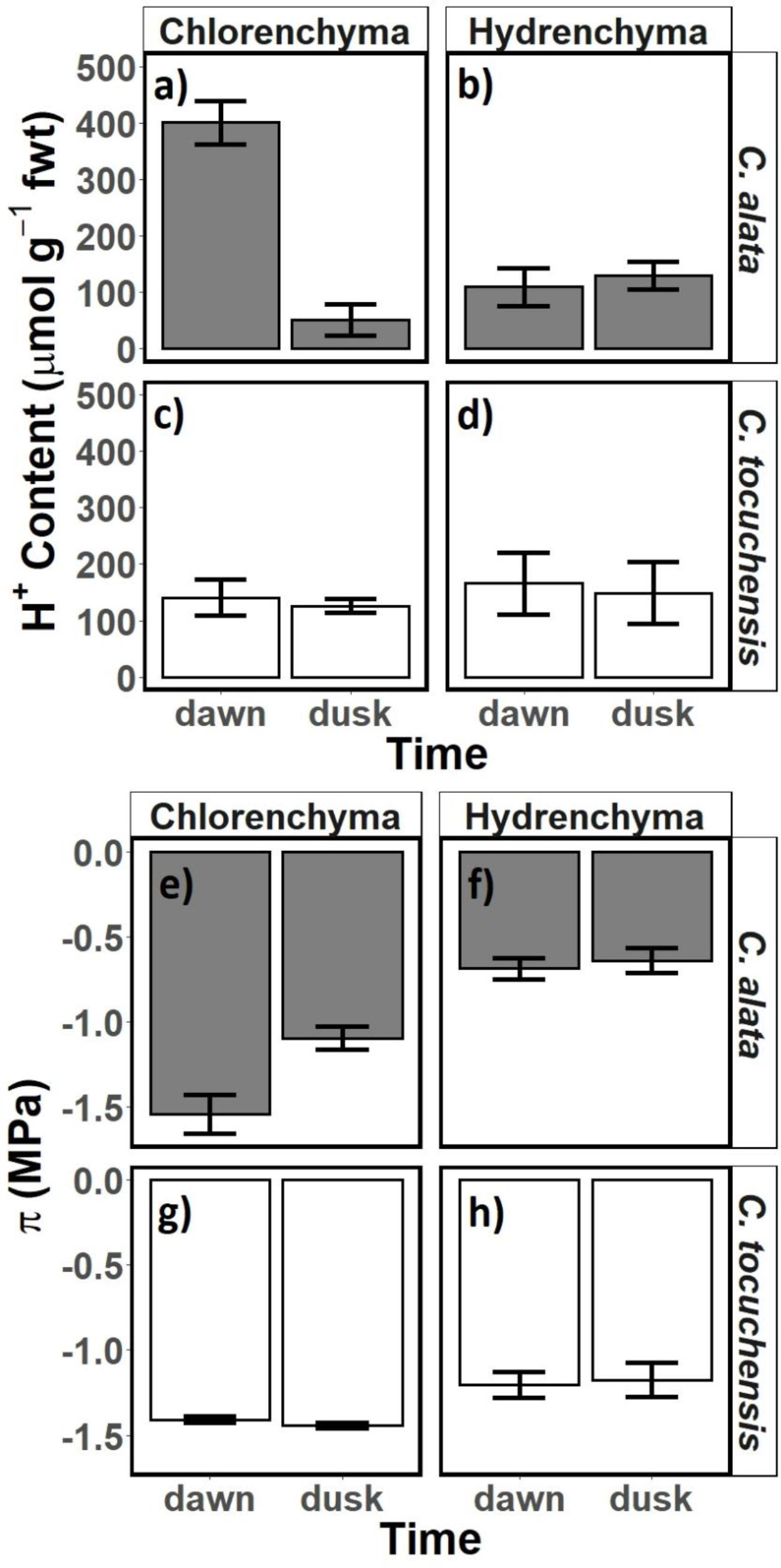
An osmotic gradient exists between hydrenchyma and chlorenchyma in *Clusia* leaves regardless of photosynthetic physiotype. (a-b) H^+^ content differs significantly between dawn and dusk in the chlorenchyma but does not differ in the hydrenchyma of the constitutive CAM species, *C. alata* (one-tailed paired t-test: p < 0.001 for chlorenchyma, p = 0.917 for hydrenchyma). (c-d) H^+^ content does not differ significantly between dawn and dusk in the chlorenchyma or between dawn and dusk in the hydrenchyma of the obligate C_3_ species, *C. tocuchensis* (one-tailed paired t-test: p = 0.136 for chlorenchyma, p = 0.142 for hydrenchyma). (e-f) Osmotic potential (π) differs significantly between dawn and dusk in the chlorenchyma but does not differ in the hydrenchyma of *C. alata*. An ANOVA showed that there is a significant difference between each measurement of osmotic potential (p < 0.001) and post-hoc Tukey- Kramer analysis found 3 significantly different groups, one represented by the chlorenchyma at dawn, another represented by the chlorenchyma at dusk and a third represented by the hydrenchyma at both dawn and dusk. (g-h) π does not differ significantly between dawn and dusk for either chlorenchyma nor hydrenchyma, in *C. tocuchensis*. However, π does differ significantly between chlorenchyma and hydrenchyma in *C. tocuchensis*. ANOVA showed that there is a significant difference between each measurement of osmotic potential (p < 0.001) and post-hoc Tukey-Kramer analysis found 2 significantly different groups, one represented by the chlorenchyma at both dawn and dusk, and another represented by the hydrenchyma at both dawn and dusk. For all graphs, n = 4 and error bars represent ± 1 standard deviation.

### Diel Changes to Hydrenchyma Relative Water Content (RWC)

We hypothesised that elevated ε would allow hydrenchyma to readily inflate and deflate, to match the hydraulic needs of the leaf. Over a 24 h period, the RWC in the hydrenchyma mirrored the stomatal conductance for well-watered *C. tocuchensis* (C_3_) and *C. alata* (CAM) (Fig. **S7**). When stomatal conductance was highest (during the day for *C. tocuchensis* and at night for *C. alata*), the RWC of the hydrenchyma dropped by approximately 10 %. During periods of stomatal closure, the RWC of the hydrenchyma was restored to 100 % (Fig. **S7**).

### CAM is More Effective Than CFT at Buffering Leaf Water Potential During Water-Deficit Stress

To disentangle the roles that CAM and C_FT_ play during drought, the Photo3 model was parameterised using physiological data included here for *Clusia*. The model allowed us to alter CAM and C_FT_ independently, and hence to assess their relative contributions to plant performance during a dry-down simulation. Four scenarios were simulated: high capacitance (HC) and low capacitance (LC) values in both a C_3_ and a CAM leaf (HC-C_3_, LC-C_3_, HC-CAM and LC- CAM, respectively). We investigated the model over the course of a diel period beginning 11 days after the cessation of rain (**Fig. 7**). Both C_3_ simulations exhibited elevated net diel photosynthetic assimilation (A_n_) and transpiration (E), in comparison to the CAM simulations (**Fig. 7**). For all simulations, the minimum daily water potential (Ψ_min_) occurred during the day. Furthermore, both CAM simulations experienced higher (less negative) Ψ_L_, which remained stable over the 24 h diel cycle unlike the C_3_ simulations.

**Fig. 7.**
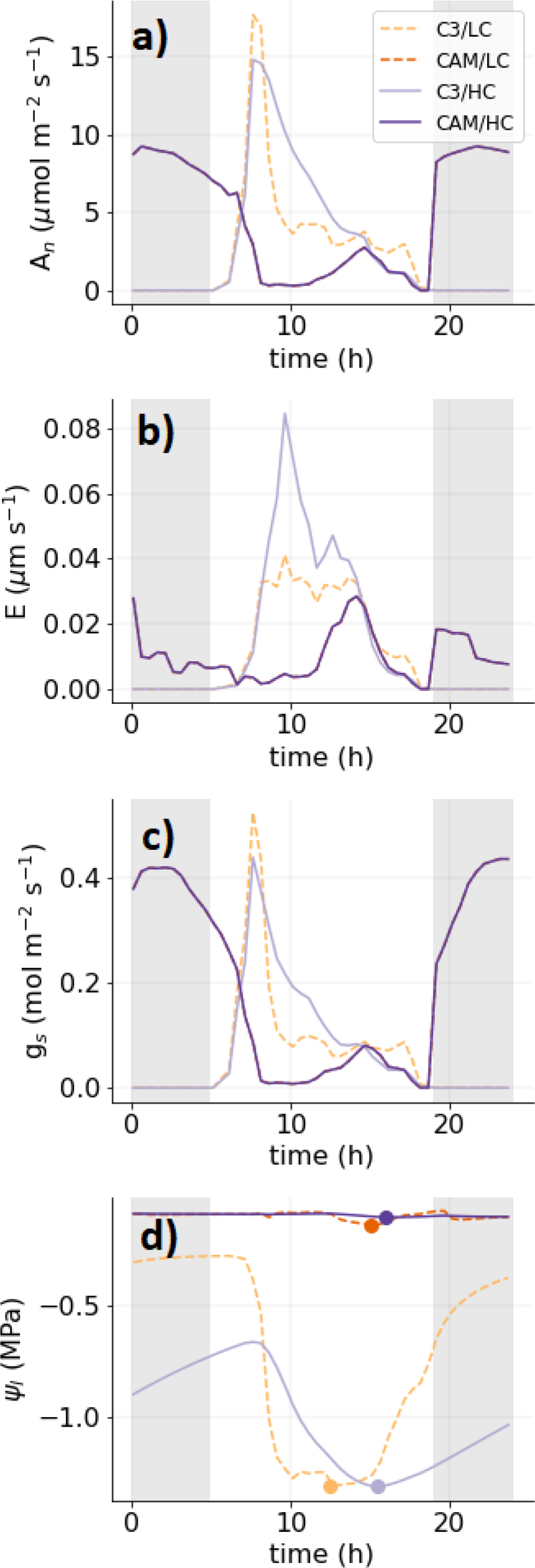
Crassulacean acid metabolism (CAM) is a more effective adaptation against drought-stress than bulk hydraulic capacitance (C_FT_) in *Clusia* leaves. All graphs show simulated diel plant physiology, estimated using the Photo3 model after 11 days of a Panamanian dry season. (a) Diel net CO_2_ assimilation (A_n_). (b) Diel transpiration (E). (c) Diel stomatal exchange of H_2_O (g_s_). (d) Diel leaf water potential (Ψ_L_).

Over a 24 h period, there was a clear difference in E and A_n_ between the HC-C_3_ and LC-C_3_ simulations; for C_3_ plants, elevated C_FT_ allowed stomatal conductance (g_s_) to remain higher for longer, during the day. In contrast, changing C_FT_ had no discernible impact in the CAM simulations. Therefore, the presence of CAM precluded any impact of C_FT_ on diel gas exchange. A similar pattern was observed when diel Ψ_L_ was analysed. The HC-C_3_ simulation showed substantially less diel fluctuation in Ψ_L_ than the LC-C_3_ simulation. However, diel patterns of Ψ_L_ were almost identical in the HC-CAM and LC-CAM simulations. When considering the simulation of a dry season drought event, elevated C_FT_ resulted in a higher (less negative) Ψ_min_ value in both C_3_ and CAM simulations. However, this only translated to a difference in cumulative E in the C_3_ background: cumulative E was higher in the HC-C_3_ than the LC-C_3_ simulation but was not substantially different between HC-CAM and LC-CAM. C_FT_ had very little impact on A_n_ or WUE (besides affecting WUE when stomata were starting to shut, and gas exchange was very low). Together, these data demonstrate that the gas exchange profile of CAM plants is more effective than C_FT_ at both minimising E and buffering Ψ_L_.

### CAM Obviates the Effect of CFT During Prolonged Drought

In addition to investigating the effect of CAM and C_FT_ over a 24-hour period, the Photo3 model was used to simulate a dry-down based on climatic data from Gamboa, Panama (**Fig. 8**), where *Clusia* plants are known to grow. Both CAM simulations experienced lower cumulative E than the C_3_ simulations. In the C_3_ simulations only, higher C_FT_ caused a small increase in cumulative E after 15 days. This supports the hypothesis that CAM is more effective than C_FT_ at mitigating transpiration losses over the dry season. Furthermore, throughout the dry-down simulation, Ψ_min_ was higher (less negative) in both CAM simulations, than in either C_3_ simulation. C_FT_ also caused a reduction in Ψ_min_ throughout the dry-down simulation for both photosynthetic types. However, CAM was substantially more effective than C_FT_ in mitigating Ψ_min_. These dynamics had a clear impact on A_n_. At the beginning of the dry-down, A_n_ was higher in both the C_3_ scenarios than in either CAM scenario. However, approximately 18 days into the dry-down, cumulative A_n_ began to plateau in both C_3_ scenarios. In contrast, both CAM simulations maintained photosynthetic assimilation for 38 days. Consequently, by the end of the dry-down, both CAM scenarios had assimilated 65% more carbon than either C_3_ scenario. In contrast, C_FT_ had very little effect on A_n_. Therefore, CAM was also more impactful than C_FT_ at maintaining net photosynthetic assimilation during drought. Due to higher A_n_ and lower E rates, both CAM scenarios exhibited higher WUE than either C_3_ scenario. In contrast, C_FT_ had little effect on WUE. In summary, CAM was substantially more effective than C_FT_ at reducing E and Ψ as well as maintaining net CO_2_ assimilation and WUE, during the drought simulation. Taken together, these analyses indicate that the presence of CAM in *Clusia* largely obviates the effect of C_FT_ over 24 h and over the course of a dry season, as neither carbon gain or water use was affected by changes to C_FT_ in CAM simulations.

**Fig. 8.**
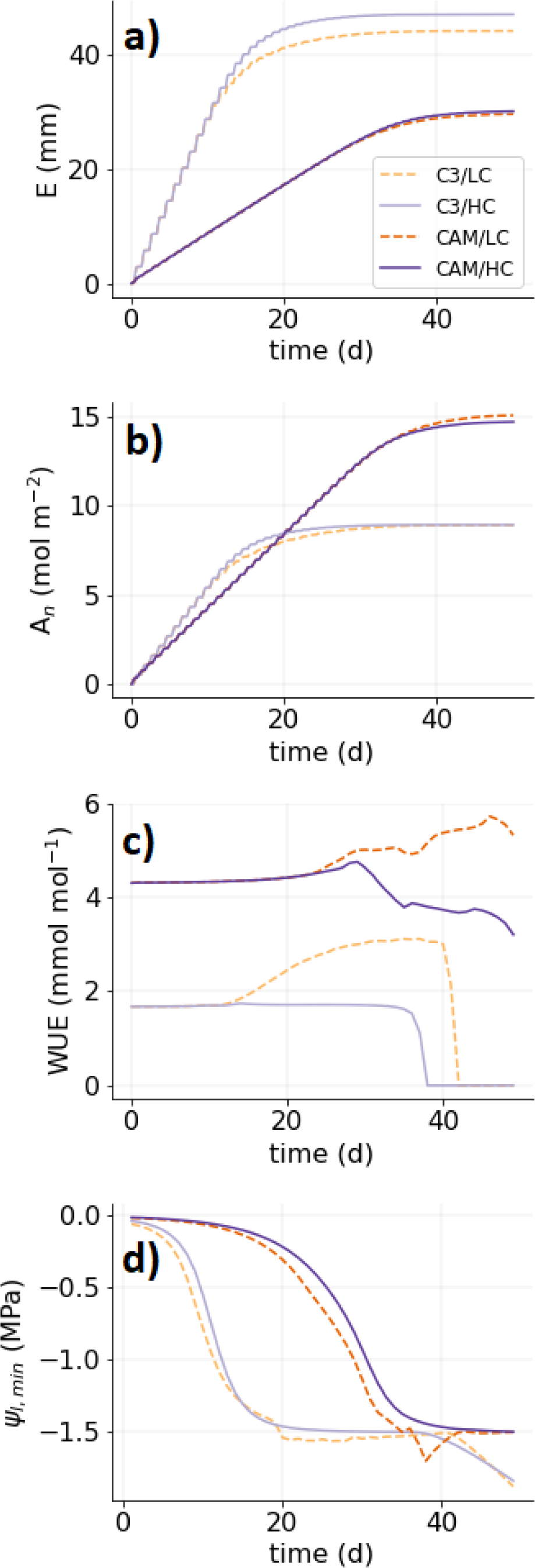
Leaf physiological traits simulated over a Panamanian dry season. (a) Cumulative transpiration (E). (b) Cumulative photosynthetic assimilation (A_n_). (c) Daily water use efficiency (WUE). (d) Daily minimum leaf water potential (Ψ_min_).

## Discussion

### Large Palisade Chlorenchyma Cells are an Adaptation for CAM, not CFT, in Clusia

Previous work on this *Clusia* collection has shown that palisade cell size and depth positively correlates with CAM photosynthesis (Barrera-Zambrano *et al.*, 2014; Borland *et al.*, 2018). This finding is congruous with observations in other taxa, where chlorenchyma cell size is greater in CAM species than C_3_ relatives (Heyduk *et al.*, 2016; Males, 2018). However, conclusions based on correlations between anatomical adaptations and CAM suffer from the potentially confounding effect of C_FT_. Moreover, it has been unclear if large chlorenchyma cells are in fact a requirement of CAM or if instead they are acting to increase C_FT_ to hydraulically buffer the leaf, as this could also act as an adaptation to drought (Edwards, 2019). In *Clusia*, we found that both CAM and chlorenchyma cell sizes were independent of C_FT_ (**Fig. 2,3**). For example, the leaves of *C. multiflora* (obligate C_3_) had C_FT_ values twice that of *C. hilariana* (constitutive CAM), despite leaf depth, palisade depth and palisade cell size being approx. 2, 3 and 4.5 times lower in the C_3_ species. *Clusia multiflora* had higher values of C_FT_ due to its investment in hydrenchyma tissue, which had a 2-fold higher tissue depth than in *C. hilariana*. The relationship between hydrenchyma and C_FT_ is also present in *Tillandsia ionantha* (Bromeliaceae), *Pyrrosia lingua* (Polypodiaceae) and *Cheilanthes myriophylla* (Pteridaceae); in which hydrenchyma tissue depth substantially buffers leaf water potentials during drought (Mcadam & Brodribb, 2013; Nowak & Martin, 1997). In *Clusia*, across all 11 species studied, we found that the hydrenchyma, and not the chlorenchyma was responsible for interspecific differences in C_FT_ (**Fig. 3**). Therefore, these analyses indicate that large palisade cells, in thick CAM leaves, are not acting to elevate C_FT_, as hydrenchyma performs this function. By accounting for the potentially confounding effect of C_FT_, we have increased the confidence that the correlation between palisade cell size and CAM is physiologically meaningful within the genus *Clusia* (Barrera-Zambrano *et al.*, 2014). Thus, this finding provides a robust demonstration that large chlorenchyma cells exist directly to aid in the storage of malic acid in the CAM cycle, rather than to hydraulically buffer Ψ_L_. This physiological framework (chlorenchyma performing CAM and hydrenchyma conferring C_FT_) provides a platform for investigations into the evolution of succulence in *Clusia*. ^13^C/^12^C isotope ratios, and hydrenchyma depth can be used as high throughput proxies for CAM and C_FT_, respectively (Leverett *et al.*, 2021; Messerschmid *et al.*, 2021). These traits could be mapped, together, onto the recently developed phylogeny of *Clusia* to understand the eco-evolutionary dynamics that have led to leaf succulence in this genus (Luján *et al.*, 2021).

We sought to understand why CAM species do not exhibit elevated C_FT_, despite having thicker leaves and more water (**Fig. 4**). Constitutive CAM species had greater WMA (**Fig. 2,4**), but these species did not exhibit elevated SWC, due to their higher LMA (**Fig. 4**). There are two possible explanations for this observation. The first is that thicker leaves are the consequence of a greater number of identical cell layers, thus elevating WMA and LMA proportionally, which would result in no change to SWC. However, this is unlikely because it is already known that CAM is associated with larger palisade cells, not more layers of identical cells, in *Clusia* (Barrera- Zambrano *et al.*, 2014; Borland *et al.*, 2018). This increases the likelihood of the second explanation: that constitutive CAM species have larger cells, whilst also investing more carbon in cell wall thickness and reinforcement. Such a scenario would cause CAM species to have elevated LMA, with no changes to SWC, despite having larger palisade cells with higher WMA values. Similar findings are observed in the genus Crassula, where larger chlorenchyma cells in thicker leaves do not amount to elevated SWC (Fradera-Soler *et al.*, 2021). This also provides a putative explanation for why the development of thick leaves in CAM species does not increase C_FT_. If large palisade cells in CAM species have thicker cell walls, they would become more rigid, which would in turn decrease C_FT_, as cell wall deformations to mobilise water would occur less readily. Chlorenchyma cell wall thickness is high in a number of CAM species, such as *Agave deserti* and *Kalanchoe daigremontiana* (Smith *et al.*, 1987; Maxwell *et al.*, 1997). Direct analysis of cell wall thickness was outside of the scope of this study, but future work should investigate the relationship between CAM and cell wall dimensions in *Clusia* and other taxa.

### Biomechanics and Osmotic Properties of Hydrenchyma Elevate CFT

Having established that the hydrenchyma, and not the chlorenchyma, is responsible for interspecific variation in C_FT_, we sought to develop a deeper mechanistic understanding of how C_FT_ is controlled in *Clusia* leaves. Across all 11 species studied, measurements of ε strongly suggest that hydrenchyma cell walls are highly elastic, and that this is responsible for their contribution to C_FT_. In addition, hemicelluloses, which are known to provide structural reinforcement to cell walls (Vanzin *et al.*, 2002; Kim *et al.*, 2020; Panter *et al.*, 2020), were found at significantly lower abundances in the hydrenchyma in comparison to the chlorenchyma (**Fig. 5**). This finding is congruent with a recent demonstration that hemicelluloses are less abundant in the leaves of phylogenetically diverse succulent species, compared to non-succulent plants (Fradera-Soler, in prep). Furthermore, pectins, which provide flexibility and can increase water holding capacity of the apoplast, were more abundant in the hydrenchyma than the chlorenchyma in *C. alata* (Fig. **S5**) (Goycoolea and Cárdenas, 2003; Sáenz *et al.*, 2004; Braybrook *et al.*, 2012; Ahl *et al.*, 2019; Fradera-Soler *et al.*, 2022). Together, these findings support the hypothesis that elastic cell walls are contributing to the elevated C_FT_ provided by the hydrenchyma. Elasticity allows cells to contort and mobilise water to meet the needs of the leaf. Cell walls are known to deform and fold in regular patterns in the hydrenchyma of *Pyrrosia* and *Aloe* and in the cortex of cacti, in order to release stored water during drought (Ong *et al.*, 1992; Mausethi, 1995; Ahl *et al.*, 2019). This allows hydrenchyma tissue to shrink as the leaf dehydrates (Fig. **S7**), thereby mitigating water loss from the chlorenchyma and maintaining Ψ_L_ at stable levels (Nowak & Martin, 1997).

Whilst elasticity allows cell walls to bend, the movement of water will not occur without an osmotic gradient between tissues. We found that both *C. alata* (constitutive CAM) and *C. tocuchensis* (obligate C_3_) maintained an osmotic gradient between the hydrenchyma and chlorenchyma (**Fig. 6**). Importantly, diel differences in chlorenchyma π, caused by CAM-related malic acid accumulation/depletion did not eradicate this osmotic gradient, meaning water can favourably move out of the hydrenchyma at all times of the day (**Fig. 6**). A similar within-leaf osmotic gradient occurs in *Agave*, and is integral to providing C_FT_ (Smith *et al.*, 1987; Schulte and Nobel, 1989). Together, highly elastic cell walls in conjunction with an osmotic gradient mean that interspecific variation in hydrenchyma depth confers a more than 4-fold variation in C_FT_, despite this tissue only comprising 7.5 – 30 % of total leaf depth.

### CAM and CFT Affect Growth and Water Loss Differently During Drought: Ecological Implications

With the advent of global warming, succulent species are becoming increasingly competitive with sympatric non-succulent plants. For example, *Mesembryanthemum crystallinum* (Aizoaceae) and *Cylindropuntia imbricate* (Cactaceae) begin to outcompete sympatric C_3_ and C_4_ grasses under drier conditions (Yu *et al.*, 2019; Huang *et al.*, 2020). In addition, succulent species have become invasive aliens in a number of ecosystems across the world (Osmond *et al.*, 2008; Novoa *et al.*, 2015; Herrando-Moraira *et al.*, 2020). Field trials are underway to utilise this adaptive advantage by growing succulent *Agave* and *Opuntia* as bioenergy crops in marginal lands (Davis *et al.*, 2017; Neupane *et al.*, 2021). However, little has been done to quantify to what extent CAM and C_FT_ are relatively responsible for any competitive advantage succulent plants experience under arid conditions. Having established that CAM and C_FT_ are independent traits in *Clusia* leaves, we investigated the roles that each trait plays during drought, by simulating Panamanian dry season conditions with the Photo3 model (Hartzell *et al.*, 2018). Our model indicated that CAM was more effective than C_FT_ at mitigating plant water stress in *Clusia* (**Fig. 7,8**). Over a 24-hour period, CAM substantially minimised fluctuations in Ψ_L_. This finding is congruent with recent work done in South Africa’s Succulent Karoo, which found that the CAM species *Malephora purpureo-crocea* (Aizoaceae) exhibited substantially dampened diel fluctuations to Ψ_L_, compared to the sympatric C_3_ species *Augea capensis* (Zygophyllaceae) (Veste & Herppich, 2021). Our model also showed that whilst elevated C_FT_ did reduce fluctuations in Ψ_L_, this effect was considerably less than that of CAM. Moreover, the presence of CAM resulted in plants losing less water from transpiration. Consequently, Ψ_min_ stayed relatively constant for about 15 days longer into the dry season in the CAM simulations, compared with the simulation of C_3_ plants (**Fig. 8**). In addition, CAM had a dramatic benefit to A_n_ during drought. Despite the CAM plants assimilating less CO_2_ than the C_3_ plants under well-watered conditions, the ability to mitigate water loss prevented drops in Ψ_L_ and allowed stomata to stay open for longer into the dry season (**Fig. 8**). Hence, by the end of the 50-day dry-down both CAM simulations had achieved 65% higher cumulative carbon uptake than either C_3_ scenario. In contrast to CAM, C_FT_ had only a small impact on gas exchange and Ψ_L_ (**Fig. 8**). Therefore, the benefits achieved from using CAM to prevent water loss appear to far outstrip those gained from using C_FT_ to buffer Ψ_L_, in *Clusia*. Together, these results indicate that CAM is more beneficial to photosynthesis and water conservation than elevated C_FT_; suggesting that this metabolic adaptation can drive the increased competitive advantage of succulent plants in nature.

In addition to comparing the contributions of different adaptations during drought, modelling can provide valuable insights into the ways in which physiological traits interact with different forms of photosynthesis (Iqbal *et al.*, 2021). Our model found that C_FT_ had a substantially smaller effect on gas exchange rates and diel Ψ_L_ in the CAM background, in comparison to the C_3_ simulations (**Fig. 7,8**). This suggests that the presence of CAM largely obviates the effect of hydrenchyma-derived C_FT_ as an adaptation to drought. These predictions agree with the recent ecological finding that CAM, and not hydrenchyma depth, correlates with environmental precipitation deficits across the ecological range of *Clusia* (Leverett *et al.*, 2021). However, the outcome of this modelling questions why hydrenchyma depth and C_FT_ vary so much across *Clusia* (Fig. **S2**). Why do species evolve deeper hydrenchyma tissue, if CAM is a more effective adaptation to drought? One hypothesis is that hydrenchyma functions more effectively as an adaptation to cloud cover at high altitudes. In high elevation tropical forests, clouds can often obscure the sun for large portions of the day (Pierce *et al.*, 2002). Being able to buffer Ψ_L_ for short periods of direct sunlight in montane cloud forests would allow leaves to maximise A_n_ when light is momentarily available, as stomata could stay open, despite water loss. An analogous phenomenon is known to occur in shade leaves; if stomata can stay open despite water loss, shade leaves will benefit from high gas exchange rates when a transitory sun fleck hits (Schymanski *et al.*, 2013). Hydrenchyma depth is often high in species living in montane cloud forests (Earnshaw *et al.*, 1987; Tanner & Kapos, 1982); and this trend held true in *Clusia*. In a pilot study of Panamanian *Clusia* species, we found that hydrenchyma depth was higher in species living at high elevation compared to those sampled in the lowlands (Supplementary File 2). More work is required to explore the ecophysiological connection between hydrenchyma depth and altitude in tropical trees. Nevertheless, it is certain that hydrenchyma, and the elevated C_FT_ this tissue provides, is a physiologically distinct adaptation, independent of CAM, in the genus *Clusia*.

## Conclusions

CAM is often found alongside tissue succulence, which has led many to speculate that this photosynthetic adaptation is always accompanied by elevated C_FT_. In this study, we showed that CAM and leaf thickness are independent of C_FT_, in the genus *Clusia*. In addition, the anatomical adaptations accompanying CAM (large palisade chlorenchyma cells) do not confer additional C_FT_, as this function is performed by the specialised hydrenchyma tissue. The lack of a relationship between chlorenchyma cell size and C_FT_ increases the likelihood that the former adaptation has evolved specifically for CAM, and not to serve any other purpose. Finally, modelling gas exchange and water relations during a Panamanian dry season predicted that CAM is substantially more beneficial than C_FT_ in buffering net carbon gain and leaf water status during drought, in *Clusia*. This highlights the benefits of CAM and demonstrates the importance of understanding this photosynthetic adaptation in a hotter, drier world. Furthermore, our physiological framework for describing the relationship between leaf anatomy, CAM and C_FT_ sets a platform to explore the eco-evolutionary dynamics that have led to the emergence of different succulent phenotypes.

## Acknowledgements

The authors thank Samuel A. Logan and Helen Martin for assistance moving *Clusia* branches and plants. This research was partially funded by Newcastle University’s R. B. Cook Scholarship and a Smithsonian Tropical Research Institute Short Term Fellowship.

## Author Contributions

AL, SH and AMB devised the research. AL, AS, PF, ARA and DCT conducted wet-lab experimental work. SH conducted Photo3 modelling. AL, KW, MG, JA and AV collected Panamanian samples. Data analysis and interpretation were done by AL, SH, KW, ARA, AS, PF, WGTW and AMB. The manuscript was written by AL, SH, KW, AS, PF, ARA, WGTW, DCT and AMB.

## Data Availability

All data and R scripts are available upon request.

## Supporting Information

The following Supporting Information is available for this article:

Fig. S1 Measured vs Modelled Leaf Hydraulic Capacitance, Using Linear Mixed Effect Model

Fig. S2 Hydrenchyma Depth Across 11 Species of *Clusia*

Fig. S3 Dissection of Hydrenchyma from Chlorenchyma in *C. alata* and *C. tocuchensis*

Fig. S4 Hydrenchyma, Palisade and Spongy Mesophyll Dimensions Across 11 species of *Clusia*

Fig. S5 Pectin Abundance

Fig. S6 Extensin (Glycoprotein) Abundance

Fig. S7 Diel Stomatal Conductance and Relative Water Content of *C. alata* and *C. tocuchensis*

Fig. S8 Model schematic of the soil-plant-atmosphere continuum

Table S1 High and low scenarios for hydraulic capacitance

Table S2 Photosynthetic parameters

Table S3 Hydraulic parameters

**<u>Methods S1</u>** Model simulations were conducted using the Photo3 model of the soil-plant-atmosphere continuum to ascertain the respective roles of CAM photosynthesis and hydraulic capacitance during drought.

**<u>File S1</u>** Pilot Study: Comparison of Hydrenchyma Depth in *Clusia* species from High-Elevation and Lowland Tropical Forest in Panama (Separate File)

**Fig. S1:**
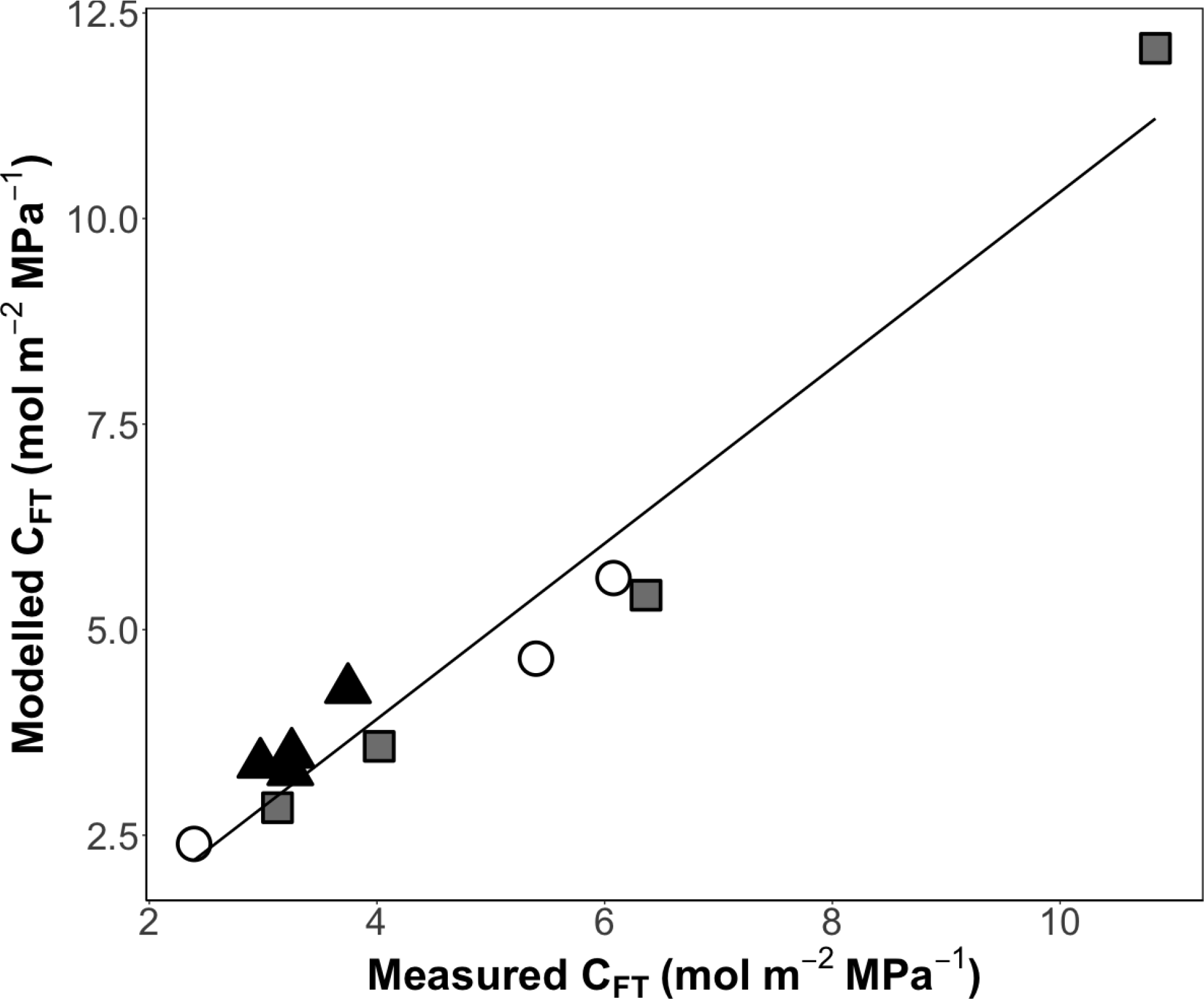
Bulk hydraulic capacitance CFT measured from pressure-volume curves is highly similar to CFT estimated from a linear mixed effect model (linear regression: slope = 1.07, R^2^ = 0.95, p < 0.0001). White circles = obligate C3, black triangles = C3-CAM intermediates, grey squares = constitutive CAM.

**Fig. S2:**
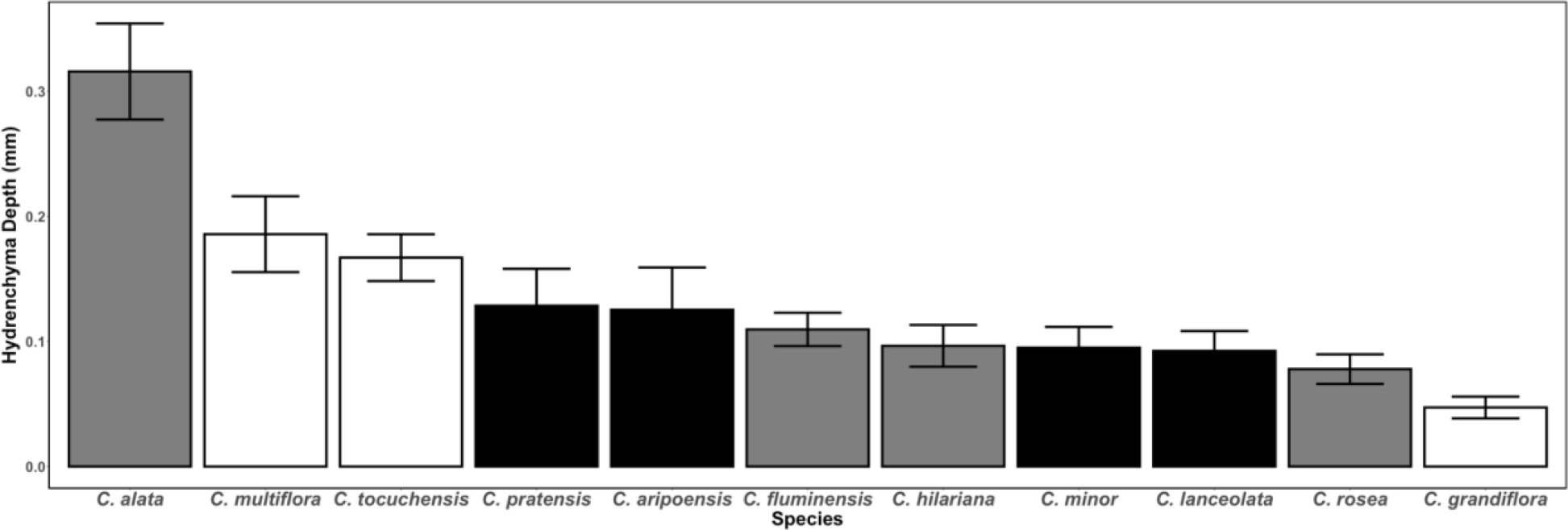
Hydrenchyma depth across 11 species of *Clusia* in the Newcastle collection. White = obligate C3, black = C3-CAM intermediate, grey = constitutive CAM; n = 12, error bars represent ± 1 standard deviation.

**Fig. S3:**
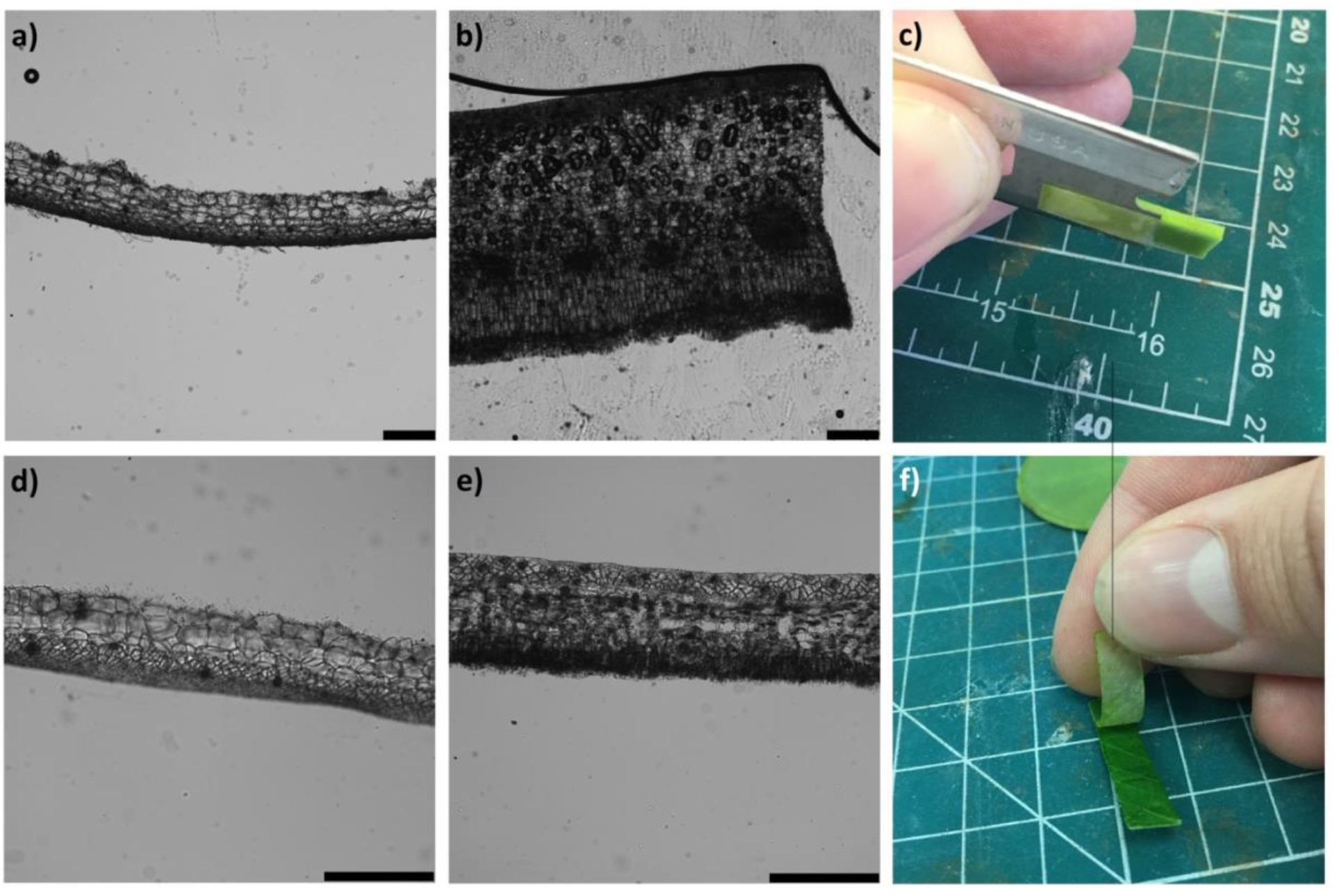
Adaxial hydrenchyma dissection from abaxial chlorenchyma. After dissecting *C. alata* the (a) hydrenchyma tissue and (b) chlorenchyma tissue are separated. Demonstration of dissection process for (c) *C. alata.* After dissecting *C. tocuchensis* the (d) hydrenchyma tissue and (e) chlorenchyma tissue are separated. Demonstration of dissection process for (f) *C. tocuchensis*. All scale bars represent 0.3 mm.

**Fig. S4:**
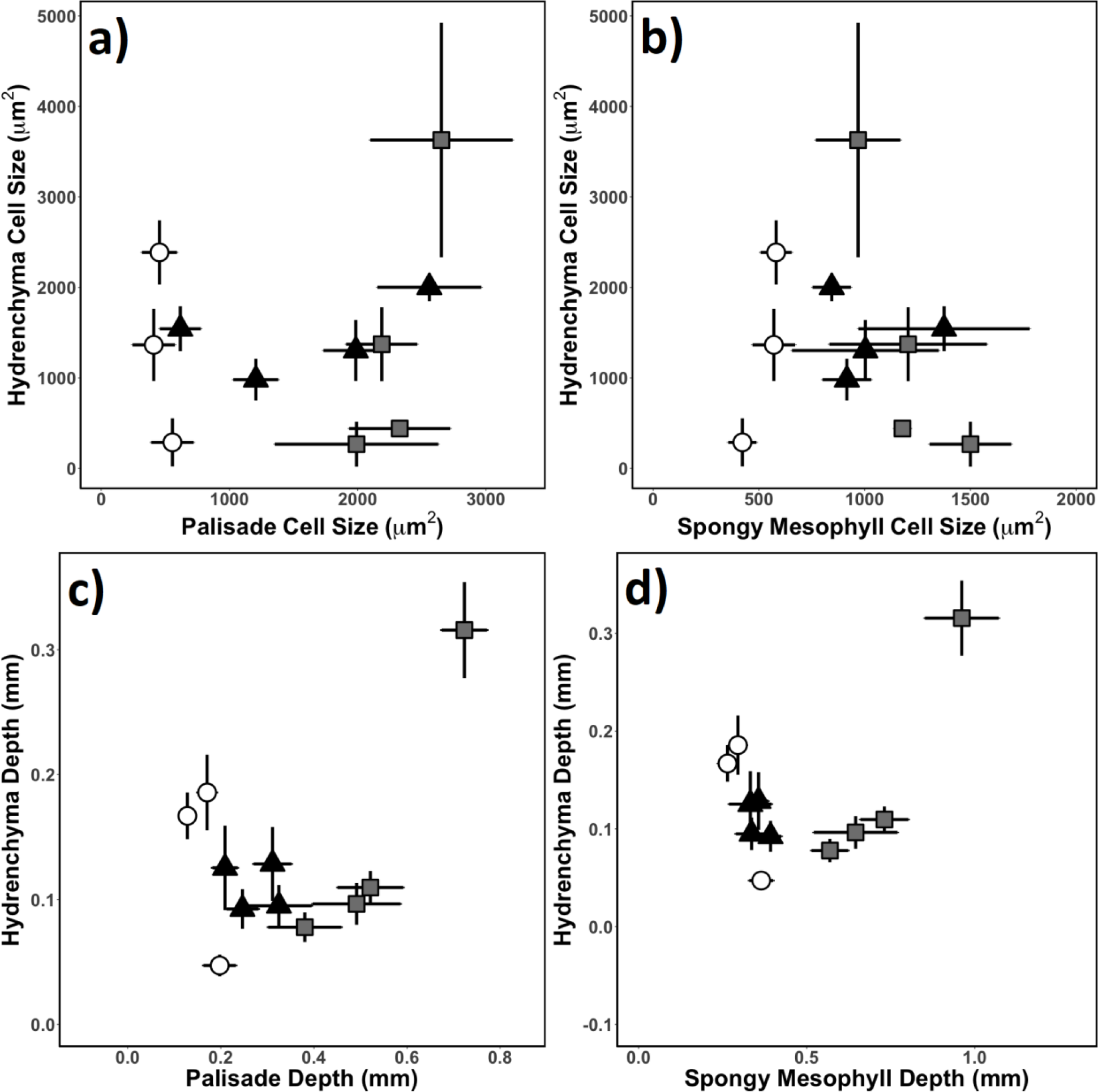
Investment in hydrenchyma is independent from investment in chlorenchyma, in *Clusia*. Across 11 species of *Clusia* there was no correlation between (a) palisade cell size and hydrenchyma cell size (p = 0.55); (b) spongy mesophyll cell size and hydrenchyma cell size (p = 0.63); (c) palisade depth and hydrenchyma depth (p = 0.16); (b) spongy mesophyll depth and hydrenchyma depth (p = 0.15). All p values are based on linear regressions. White circles = obligate C3, black triangles = C3-CAM intermediates, grey squares = constitutive CAM and error bars represent ± 1 standard deviation.

**Fig. S5:**
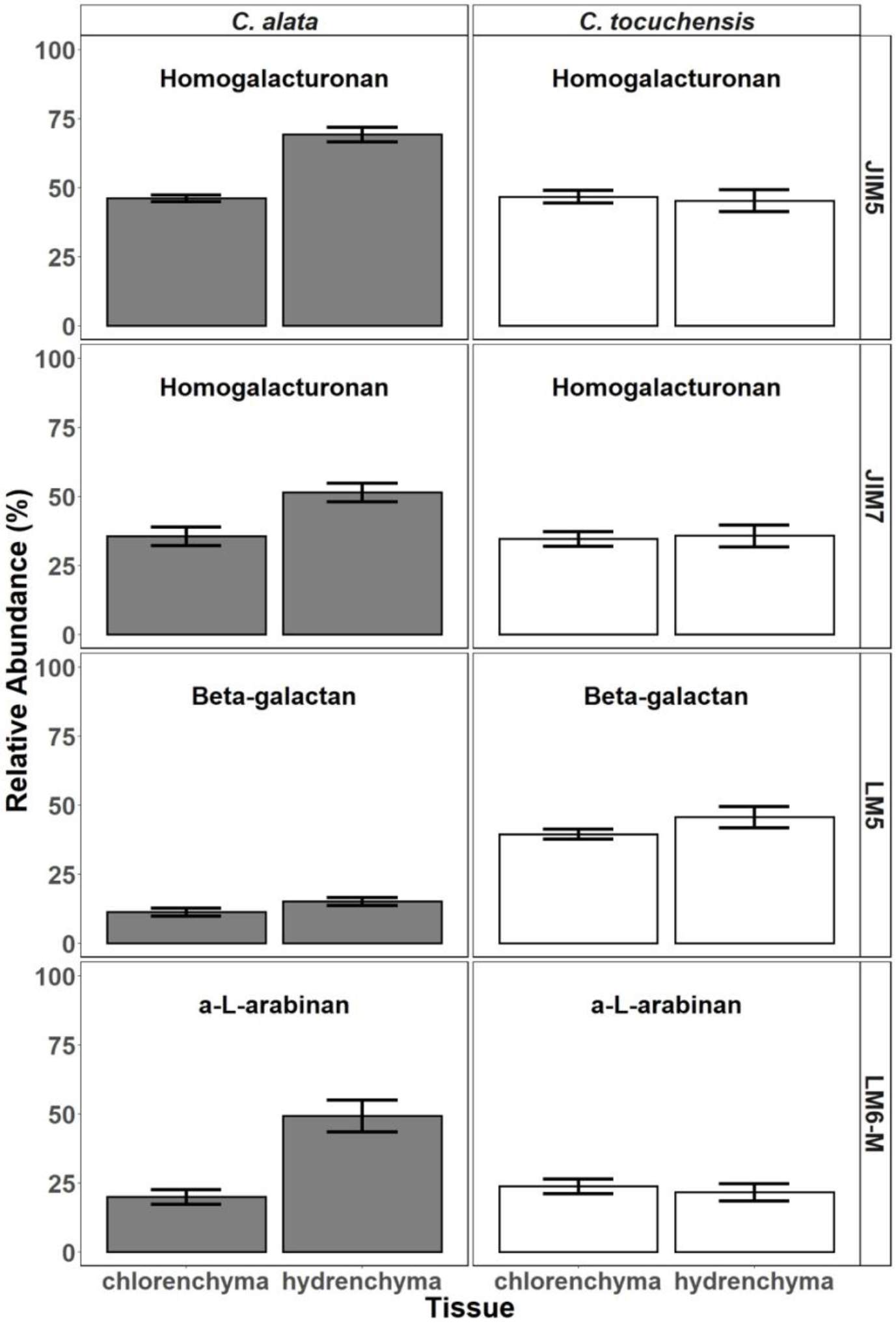
Pectin abundance of hydrenchyma and chlorenchyma in *C. alata* (constitutive CAM) and *C. tocuchensis* (obligate C3).

**Fig. S6:**
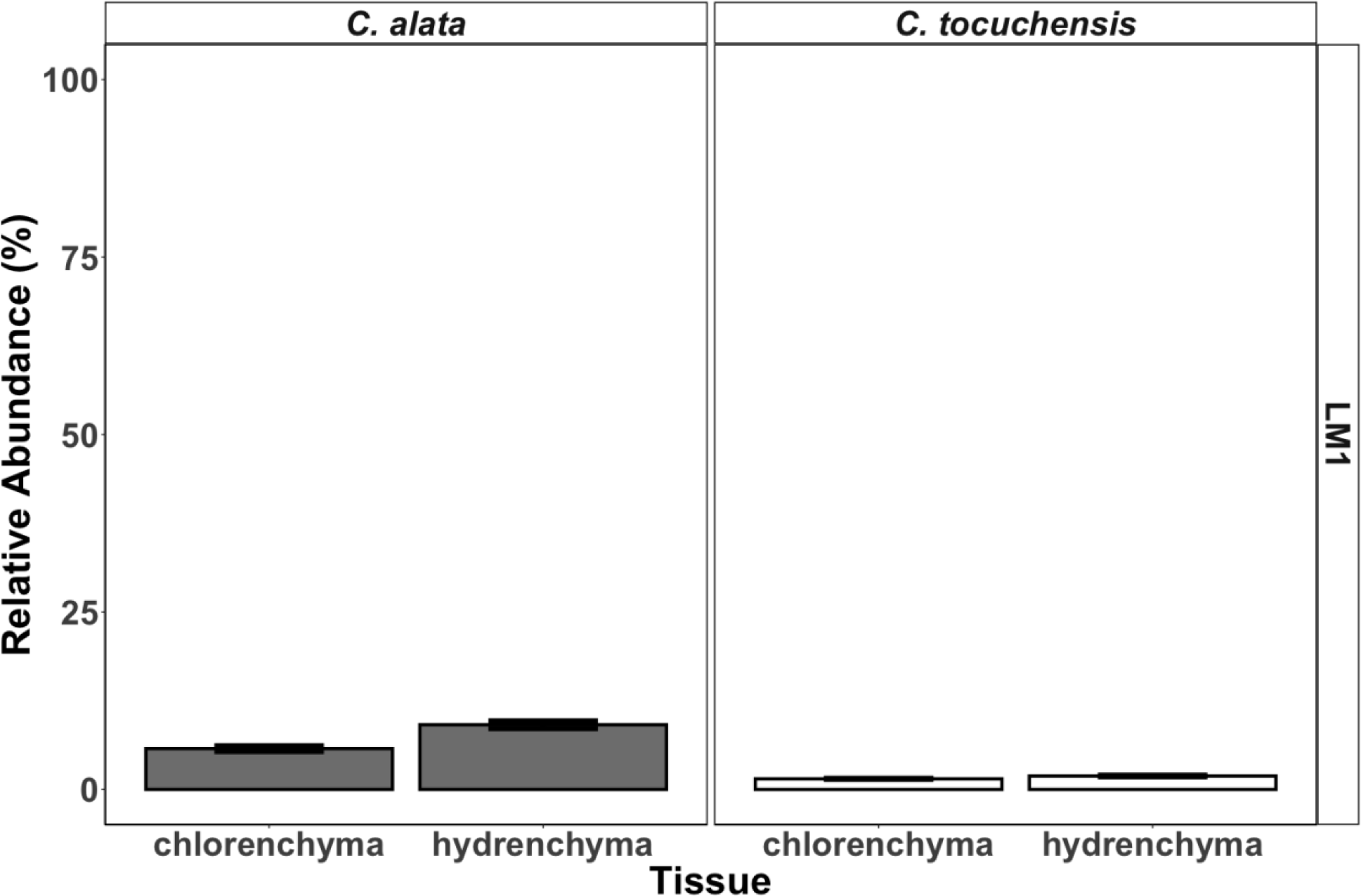
Abundance of the glycoprotein, extension in hydrenchyma and chlorenchyma tissue from in *C. alata* (constitutive CAM) and *C. tocuchensis* (obligate C3).

**Fig. S7:**
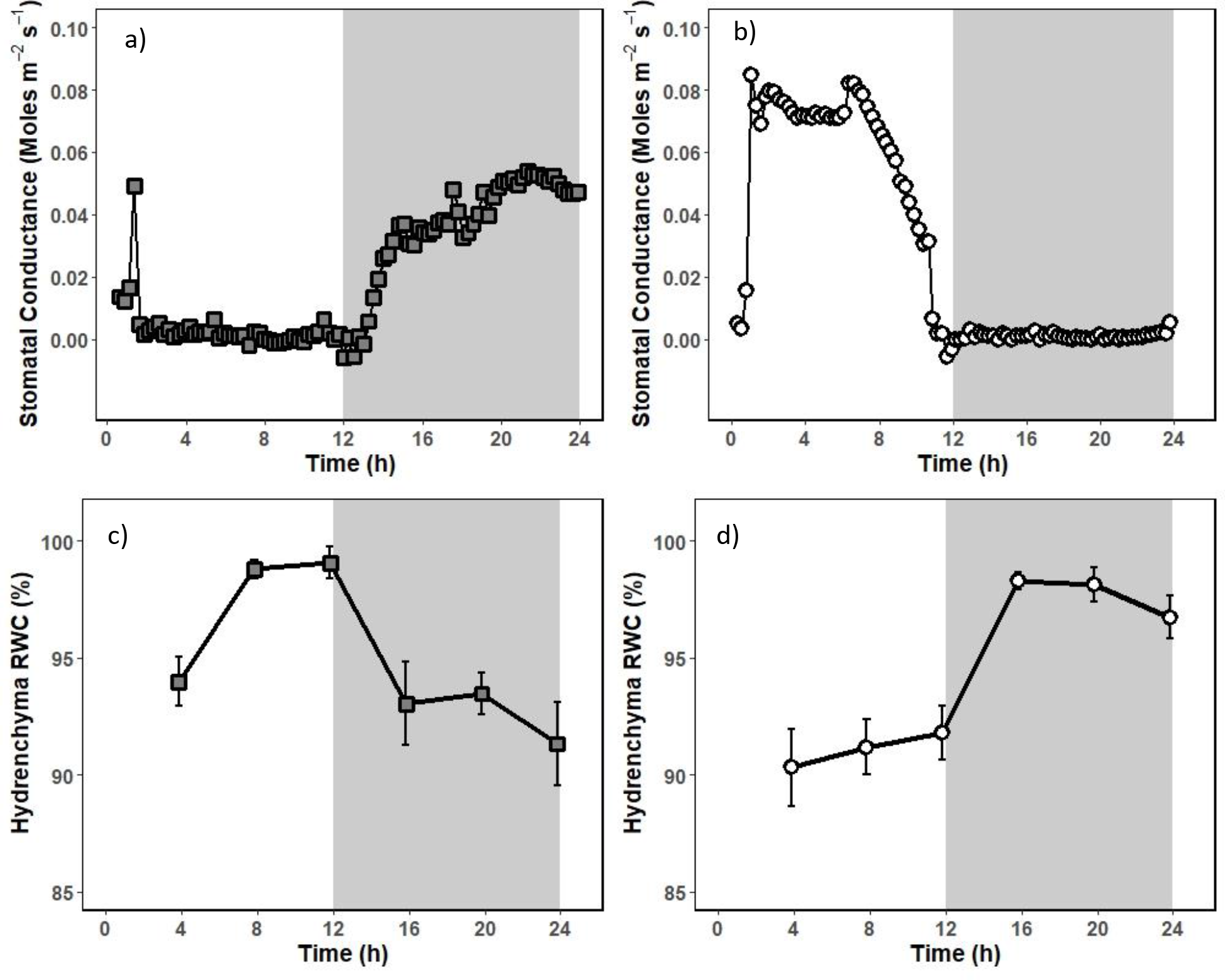
Diel changes in stomatal conductance mirror diel fluctuations in hydrenchyma relative water content (RWC). White background represent day, and shaded backgrounds represent night. Stomatal conductance was measured every 15 minutes for 24 h, in (a) the constitutive CAM species, *C. alata*, and (b) the obligate C3 species, *C. tocuchensis*. Hydrenchyma RWC was estimated every 4 hours for 24 h in (c) the constitutive CAM species, *C. alata*, and (d) the obligate C3 species, *C. tocuchensis*. For both species, RWC dropped when stomatal conductance was highest. For gas exchange, 3 replicates were recorded per species, and one representative graph is shown. For RWC data, n = 4 and error bars represent ± 1 standard deviation.

**Fig. S8:**
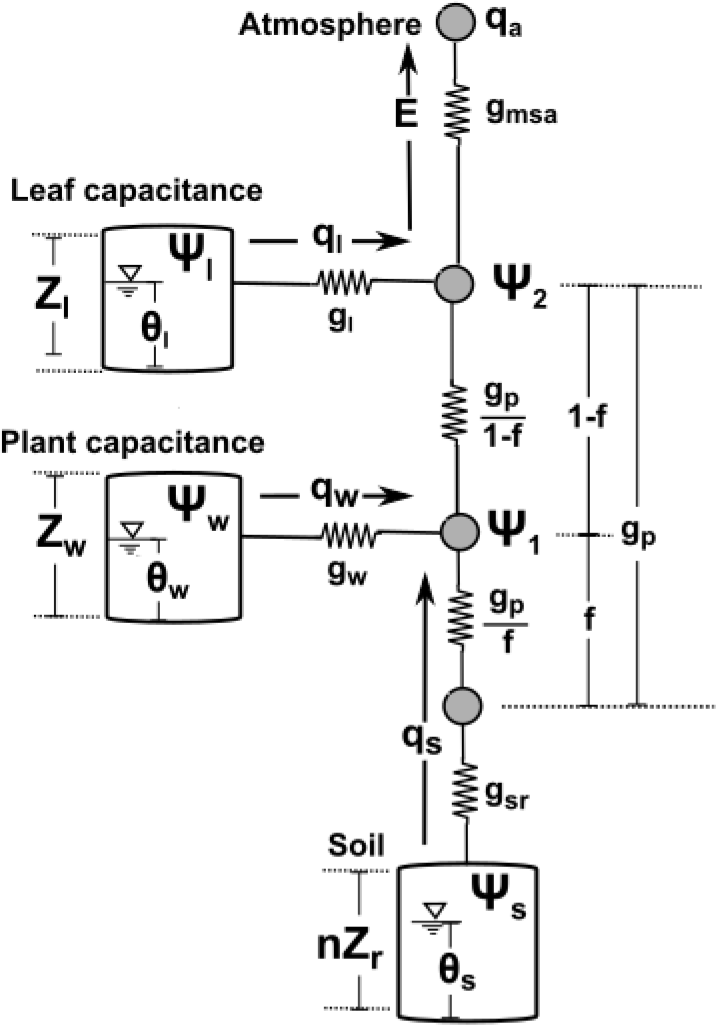
Model schematic of the soil-plant-atmosphere continuum. Soil water storage, plant sapwood capacitance (*Zw*) and leaf hydraulic capacitance (*Zl*) are accounted for and are described as capacitors in the mathematical scheme. The capacitors and connection nodes are linked through resistors, and the flux through each resistor is equal to the product of the conductance (*g*) and the water potential gradient (*ψ*).

**Table S1:** High and low model scenarios for hydraulic capacitance

| Parameter | LC scenario | HC scenario | Units | Description |
| --- | --- | --- | --- | --- |
| $c_l$ | 0.047 | 0.15 | $\text{MPa}^{-1}$ | Relative leaf hydraulic capacitance |
| WMA | 450 | 1600 | $\text{g m}^{-2}$ | Water mass per unit leaf area |
| $C_{FT}$ | 1.18<br>*(Hydrenchyma depth = 0.38 mm)* | 13.3<br>*(Hydrenchyma depth = 0.04 mm)* | $\text{mol m}^{-2}$<br>$\text{MPa}^{-1}$ | Bulk leaf hydraulic capacitance |

**Table S2.**

| Parameter | C3 Value | CAM value | Units | Description |
| --- | --- | --- | --- | --- |
| LAI | 2<sup>a</sup> | 2<sup>a</sup> | - | Leaf area index |
| V<sub>c,max0</sub> | 48.51<sup>b</sup> | 38.43<sup>b</sup> | μmol m<sup>-2</sup> s<sup>-1</sup> | Max carboxylation capacity |
| J<sub>max0</sub> | 97.01<sup>c</sup> | 76.85<sup>c</sup> | μmol m<sup>-2</sup> s<sup>-1</sup> | Max. electron transport rate |
| Ψ<sub>l0</sub> | -1.5<sup>a</sup> | -1.5<sup>a</sup> | MPa | Point of maximum plant water stress |
| Ψ<sub>l1</sub> | -0.5<sup>a</sup> | -0.5<sup>a</sup> | MPa | Onset of plant water stress |
| M<sub>max</sub> | - | 400<sup>d</sup> | mol m<sup>-3</sup> | Max malic acid storage |
| A<sub>m,max</sub> | - | 15<sup>d</sup> | μmol m<sup>-2</sup> s<sup>-1</sup> | Max rate of malic acid storage flux |

<sup>a</sup>Assumed following Bartlett, Vico, and Porporato (2014)
<sup>b</sup>Assumed to be half of Jmax following Kattge and Knorr (2007)
<sup>c</sup>Measured by Alistair Leverett (unpublished data)
<sup>d</sup>Assumed following (Taybi, Nimmo, and Borland 2004)

**Table S3:** Hydraulic parameters

| Parameter | Value | Units | Description |
| --- | --- | --- | --- |
| TLP<sub>ψ</sub> | -1.5<sup>a</sup> | MPa | Water Potential of the Turgor Loss Point |
| Z<sub>r</sub> | 0.4 | m | Rooting depth |
| g<sub>a</sub> | 95<sup>b</sup> | mm s<sup>-1</sup> | Atmospheric conductance, per unit ground area |
| g<sub>leaf</sub> | 0.1<sup>b</sup> | μm MPa<sup>-1</sup> s<sup>-1</sup> | Leaf hydraulic conductance, per unit leaf area |
| g<sub>p,max</sub> | 0.125<sup>c</sup> | μm MPa<sup>-1</sup> s<sup>-1</sup> | Xylem hydraulic conductance, per unit leaf area |
| g<sub>w,max</sub> | 0.01 | μm MPa<sup>-1</sup> s<sup>-1</sup> | Stem storage hydraulic conductance, per unit leaf area |
| vwt | 0.58<sup>b</sup> | mm | Stem water storage, per unit leaf area |

<sup>a</sup>Assumed following Leverett et al. (2021)
<sup>b</sup>Assumed following Bartlett, Vico, and Porporato (2014)
<sup>a</sup>Value for *Clusia minor* from Zotz et al. (1997) for a stem diameter of 15 mm assuming a plant height of 4 m (Borland et al. 1996)

## Methods S1

In order to explore the effects of photosynthetic pathway (C3 vs CAM), and hydraulic traits (specifically bulk leaf hydraulic capacitance) on performance in *Clusia*, the Photo3 model (Hartzell, Bartlett, and Porporato 2018) was adjusted to include leaf hydraulic capacitance in addition to stem hydraulic capacitance. The model was run using parameterisations for four hypothetical scenarios: C3 with low hydraulic capacitance, C3 with high hydraulic capacitance, CAM with low hydraulic capacitance, and CAM with high hydraulic capacitance (see Table 1). The model was run for a 60-day drydown, starting with well-watered conditions (volumetric soil moisture = 0.4, stem water content = 98%, leaf water content = 98%). Hydraulic parameter ranges were estimated from experimental data (hydraulic capacitance and water mass per area were varied concurrently in order to estimate maximum potential effect of hydraulic parameters relating to water storage). A turgor loss point (TLPΨ) of -1.5 MPa was assumed (Leverett *et al.*, 2021).

The Photo3 model combines representation of the C3, C4, and CAM photosynthetic pathway with a hydraulic description of the soil-plant-atmosphere continuum. Here we depict stem and leaf water storage through a resistor capacitor schematic, similarly to Jones (2013) and Nobel and Jordan (1983), in order to estimate the effects of leaf hydraulic properties on water use efficiency and transpiration.

The photosynthetic carbon demand by the Calvin cycle, *A*_*d*_, is given according to Farquhar, von Caemmerer, and Berry (1980) and is modified to account for effects of plant water stress, i.e.,

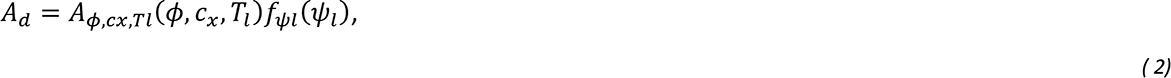

where *A*_*ϕ,cx,Tl*_(*ϕ, cx, Tl*) is the Farquhar photosynthetic demand, *ϕ* is the incoming photosynthetically active radiation, *c*_*x*_ is the internal CO_2_ concentration, and *T*_*l*_ is the leaf temperature. *fψ*_*l*_(ψ_*l*_) describes the effect of leaf water potential on photosynthetic demand and is a function of the leaf water potential, ψ_*l*_. This nonlinear behavior, with little control on photosynthetic demand at relatively high water potentials, is captured through a piecewise function (e.g. Daly, Porporato, and Rodriguez-Iturbe (2004)),

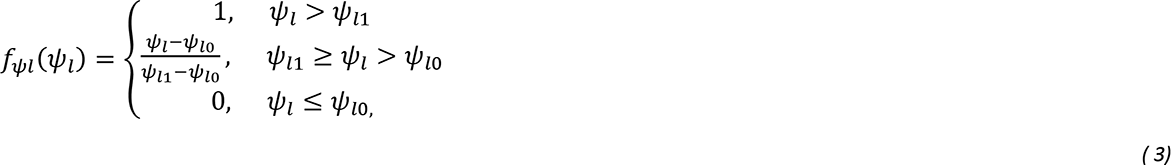

where ψ_*l1*_ is the leaf water potential at the onset of water stress and ψ_*l0*_ is the leaf water potential at which carbon assimilation approaches zero.

The net carbon uptake through the stomata, *A*_*n*_, is given by the photosynthetic carbon demand *A*_*d*_ in the case of C3 photosynthesis. In the case of CAM photosynthesis, the carbon assimilation is modified by a circadian rhythm oscillator describing the circadian state and the amount of malic acid storage (see Bartlett, Vico, and Porporato (2014) and Hartzell, Bartlett, and Porporato (2018).

The stomatal conductance to CO2, *g*_*s*_, is linearly related to the net carbon assimilation *A*_*n*_ and is modified by the vapor pressure deficit as supported by optimality approaches and empirical data (Katul, Palmroth, and Oren 2009; Hartzell, Bartlett, and Porporato 2018), i.e.

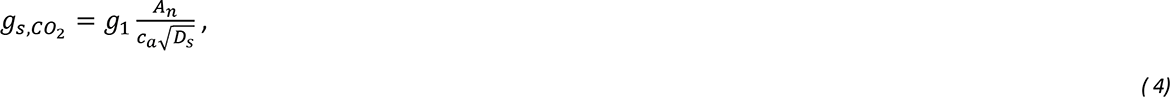

where *c*_*a*_ is the atmospheric CO_2_ concentration, *D*_*s*_ is the vapor pressure deficit, and *g*_1_ is a proportionality constant specific to the type of photosynthesis, scaled based on typical ci/ca ratios.

As simulations are conducted in the range where leaf water potential is above the TLPΨ, the leaf hydraulic capacitance is assumed to be constant in this range and the leaf water potential is assumed to be a linear function of the relative leaf water content θ_*l*_, i.e.,

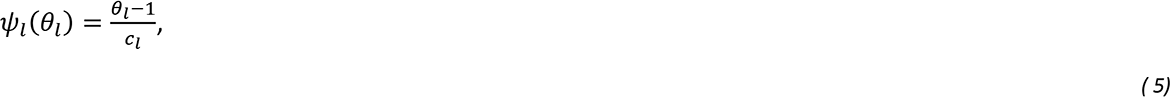

where *c*_*l*_ is the leaf-specific hydraulic capacitance before the TLPΨ. In general, fluxes between each of the storage connection nodes are given by the conductance between the nodes multiplied by the difference in water potential between the two nodes, i.e., 

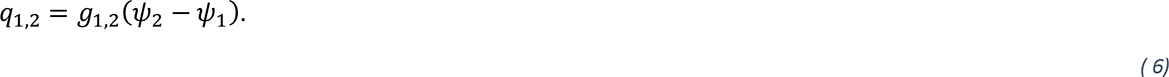

The transpiration rate is given by the difference in specific humidity between connection node 2 (*q*_2_) and the atmosphere (*q*_*a*_), multiplied by the stomatal conductance to water, which is 1.6 times the stomatal conductance to CO2 given in Equation *(3)*.

The four unknowns, i.e. the water potentials at the two connection nodes, Ψ_1_, and Ψ_2_; the transpiration rate, and the leaf temperature are solved by combining equations for the leaf energy balance, transpiration rate, and the summations of the hydraulic fluxes in and out of each of the connection nodes similarly to the procedure described in Hartzell, Bartlett, and Porporato (2017).

## Pilot Study – Possible Relationship Between High Altitudes and Hydrenchyma Depth, in *Clusia*

CAM is known to be less frequent at higher altitudes in *Clusia*, and strong CAM has not been observed in species living > 680 m above sea level (Holtum *et al*., 2004). We hypothesised that at high altitudes, species would invest in deeper hydrenchyma tissue; as CAM is a less effective adaptation to precipitation deficits in these ecosystems. We conducted a pilot study to assess differences in hydrenchyma depth between species living at high and low altitude ecosystems, in Panama. We sampled leaves from mature trees living in Cerro Jefe montane forest (1007m above sea level; 0913794N, 07922995W), and around Gamboa lowland tropical dry forest at (38m above sea level; 9.120085N, 79.701894W). Species sampled in Cerro Jefe were *C. coclensis* Standl., *C. cretosa* Hammel ined., *C. liesneri* Maguire, *C. multiflora* Kunth-complex and *C. osseocarpa* Maguire. For species sampled from Cero Jefe, n = 4-5. Species sampled in Gamboa were the C3 species: *C. cupulata* Maguire; the C3-CAM intermediate species: *C. quadrangula* Bartlett, *C. valerioi* Standl., *C. peninsulae* Hammel, *C. fructinangusta* Cuatrec., *C. pratensis* Seem., *C. minor* L., *C. uvitana* Pittier; and the constitutive CAM species *C. rosea* Jacq. For species sampled from Gamboa, n = 2-4, except for *C. fructiangusta*, for which only one individual was available. The photosynthetic phenotype of these species was determined according to Holtum *et al*. (2004).

To assess the depth of hydrenchyma in species living in the two sites, branches were cut, placed in plastic bags and returned to the laboratory at the Earl S. Tupper Centre, Panama City, Panama. The third leaf from the apex of each branch was then cut, underwater, and the cut petiole remained submerged in water to incubate. Rehydrating leaves were then placed in a box overnight, which was filled with damp tissues to increase humidity. The following day, leaves were hand sectioned, with a razor blade, and sections were imaged with a fluorescent microscope (Nikon Eclipse 600 Microscope with a Nikon Dri- 1 camera). Hydrenchyma depth was measured using ImageJ (NIH).

We found that species sampled around Gamboa had hydrenchyma depths ranging from 0.06 mm (*C. valerioi*) to 0.17 mm (*C. cupulata*). Species sampled from Cero Jefe had hydrenchyma depths ranging from 0.13 mm (*C. cretosa*) to 0.41 mm (*C. osseocarpa*). Overall, these data strongly suggest that hydrenchyma depth is greater in species living in high altitude ecosystems.

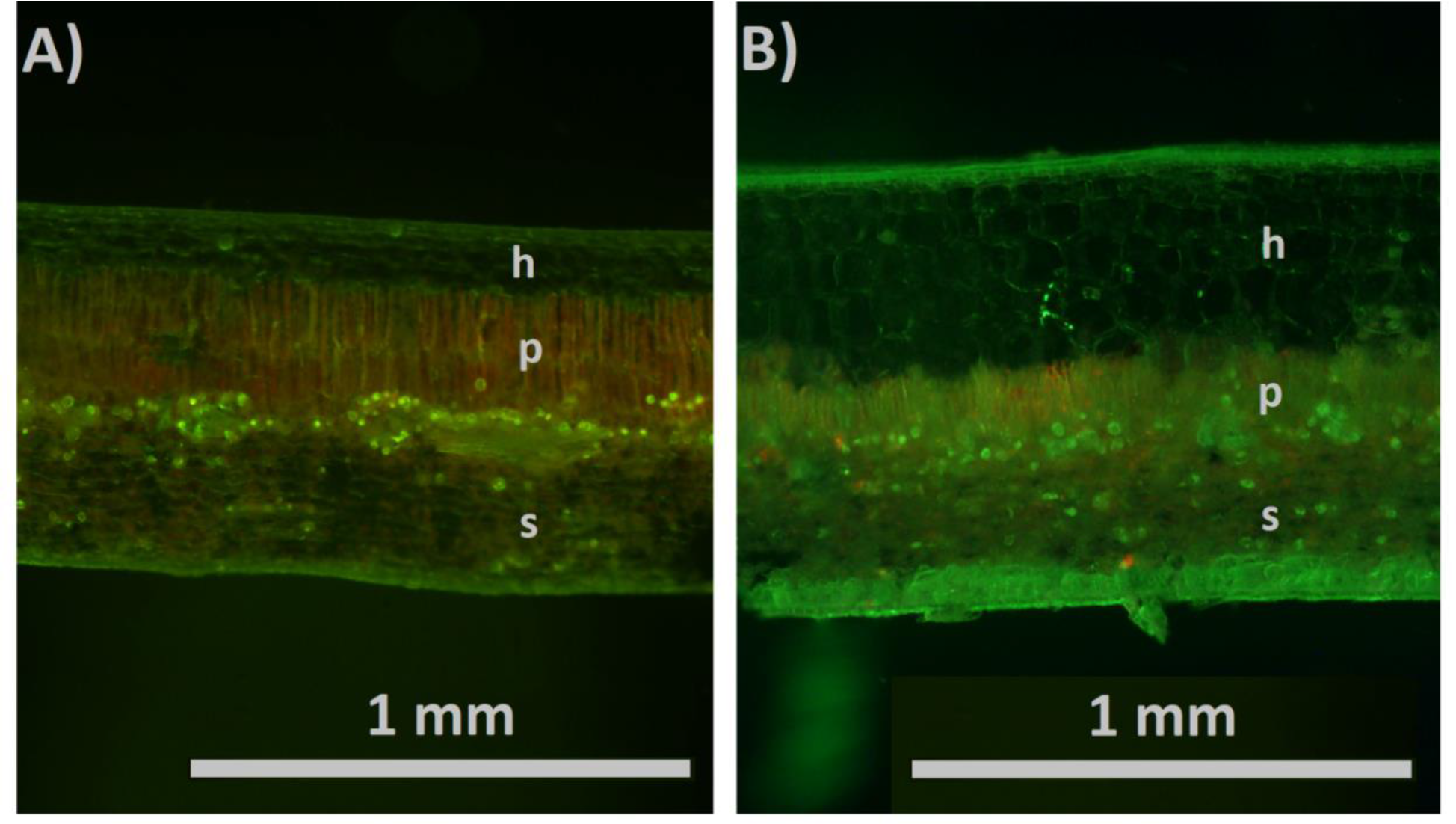

Examples of hand-sectioned leaves from field-grown *Clusia* trees. (*A*) *Clusia pratensis*, sampled in Gamboa Lowland Tropical Forest. (*B*) *Clusia osseocarpa*, sampled from Cerro Jefe montane forest. Each tissue layer is labelled: h = hydrenchyma, p = palisade, s = spongy mesophyll.

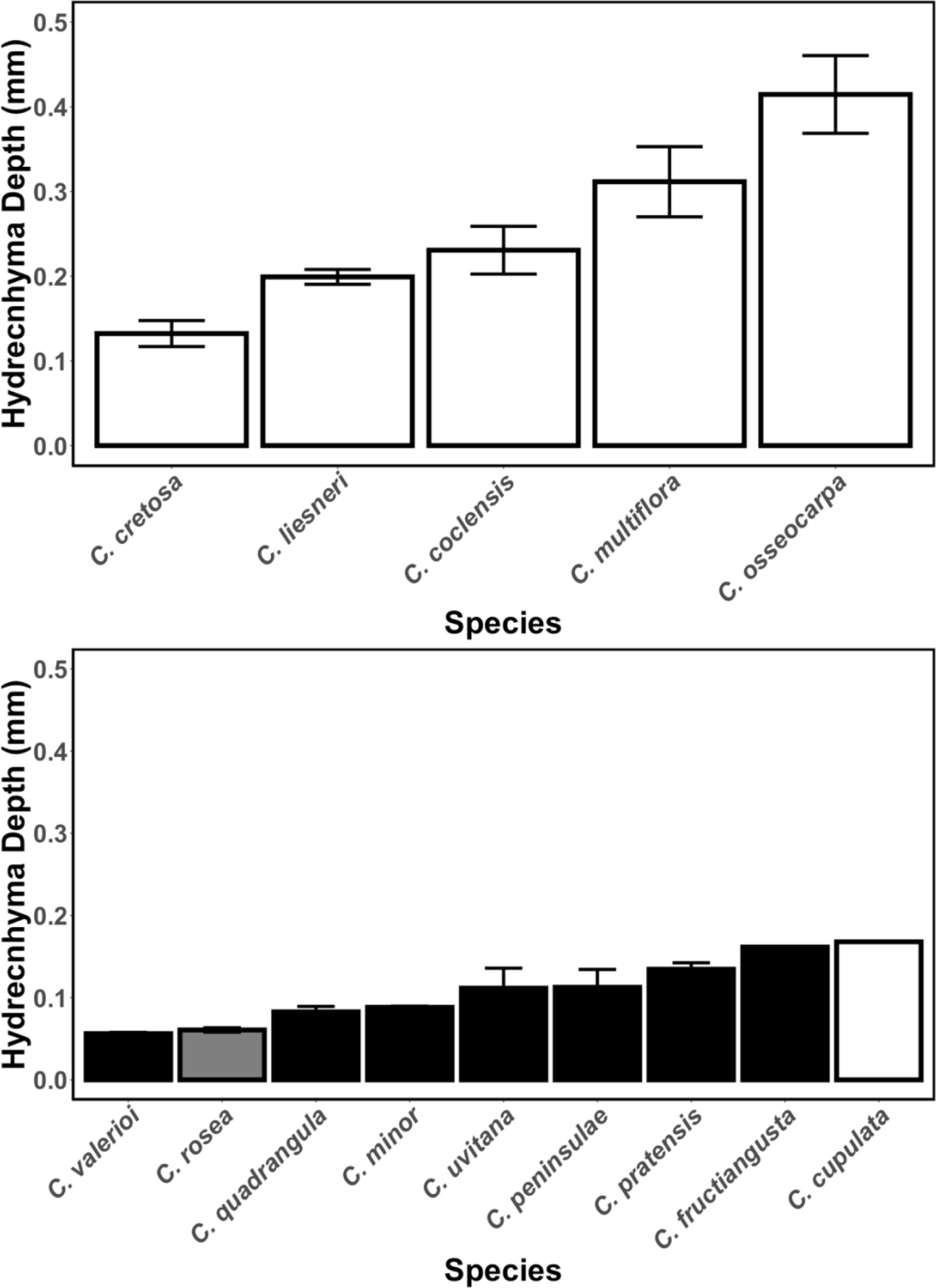

Species in cloud forest ecosystem have deeper hydrenchyma tissue than those living in lowland seasonally dry forest in Panama. The 5 C3 species found in (*A*) Cerro Jefe montane ecosystem all had deeper hydrenchyma tissue than any of the 9 species growing in (*B*) the lowland tropical forest environment in Gamboa. Error bars represent ± one SD of the mean for each species. White = Obligate C3; black = C3-CAM intermediate; grey = constitutive CAM.

